# Multimodal profiling reveals *Mycobacterium tuberculosis* restricts lung mitochondrial immunometabolism to promote pathogenesis

**DOI:** 10.64898/2026.04.07.717012

**Authors:** Hedwin Kitdorlang Dkhar, Prashant Bajpai, Ana Beatriz Enriquez, Louis Brown Hopkins, Stanzin Dawa, Jonathan Kevin Sia, Ayana Paul, Ranjna Madan-Lala, Mark Cole Keenum, AshishKumar Sharma, Adam Nicolas Pelletier, Shu Ling Goh, Terra Laretta Brown Riddick, Ted Joseph Whitworth, Keith Eric Prater, Ricardo Cèsar Guerrero-Ferreira, Jeffrey Collins, Jyothi Rengarajan

## Abstract

Early events in the lung that shape protective immune responses to *M. tuberculosis* (Mtb) infection are not well understood but are critical for developing better vaccines and immunomodulatory therapies for tuberculosis. Here, we used high-dimensional flow cytometry, single-cell transcriptomics, and untargeted metabolomics to define the early lung immune environment that precedes the development of protective versus pathogenic outcomes following aerosolized Mtb infection of mice. We show that Mtb induced sustained glycolysis in the lung while restricting oxidative phosphorylation (OXPHOS) and impairing mitochondria, in part through the Mtb serine protease Hip1, leading to low energy output and suboptimal macrophage-T cell interactions that promoted pathogenic immunity. However, robust induction of mitochondrial OXPHOS, amino acid metabolism, and fatty acid oxidation in the early lung resulted in high ATP output and enhanced innate-T cell signaling networks that stimulated protective immune responses. Moreover, we identified a novel mitochondrial immunometabolic lung signature associated with protective outcomes to Mtb infection in animal models and humans. Our studies identify induction of mitochondrial dysfunction as a mechanism employed by Mtb to manipulate lung immunometabolism to its benefit and reveal that maintenance of intact mitochondrial metabolism in the early lung is pivotal for generating protective outcomes to Mtb infection.

## Introduction

Tuberculosis (TB) is primarily a disease of the lung, and transmission of TB occurs *via* aerosol particles containing *Mycobacterium tuberculosis* (Mtb) derived from disrupted granulomatous lesions in the lungs of a person with active TB (ATB) disease^1, 2, 3, 4^. Notably, the majority of healthy individuals exposed to Mtb mount immune responses that protect against developing ATB, leading to long-term asymptomatic infection^5, 6^. However, in ∼5% of infected individuals who develop primary TB disease, pathogenic immune responses promote destructive lung pathology and fail to control bacterial replication^7, 8, 9, 10^. Early events in the lung drive the balance between subsequent protective *versus* pathogenic immune responses and are critical for shaping the outcome of Mtb infection. Interactions between Mtb and tissue-resident alveolar macrophages (AMs), followed by interstitial macrophages (IM) and monocyte-derived macrophage responses^11, 12, 13, 14^, lead to the recruitment of other innate immune cells, including neutrophils, and natural killer (NK) cells^15, 16, 17^. However, Mtb enables bacterial replication within permissive macrophages in the lung, impairs host antimicrobial responses, and impedes dendritic cell (DC) functions through multiple virulence factors^4, 18^. Animal models of TB susceptibility have shown that these early innate immune evasion strategies lead to delayed recruitment of T cells into the lung that fail to eradicate Mtb and instead result in lung pathology^15, 19^. Mtb-specific effector CD4^+^ T helper-1 (Th_1_) cells producing cytokines such as IFN-y and TNF-α migrate into the lung parenchyma and are necessary but not sufficient for conferring protective immunity against TB^16, 20, 21, 22^. While Th_17_ responses in the lung are associated with protective outcomes during Mtb infection^23, 24^ or vaccination^25, 26^, we have shown that Mtb restricts Th_17_ polarization by preventing optimal innate-T cell crosstalk ^27, 28^. However, the mechanisms governing early immune events in the Mtb-infected lung that shape pathogenic *versus* protective immune outcomes remain poorly understood. We have previously established that the immunomodulatory serine protease Hip1 (Rv2224c) is a critical Mtb virulence factor that impairs host innate and adaptive immunity, thereby promoting lung immunopathology. Hip1-mediated proteolysis of its substrate GroEL2 dampens macrophage activation and proinflammatory cytokine production and restricts lung Th_17_ responses by impeding DC-T cell costimulatory pathways^27, 29, 30^. In the absence of Hip1, Mtb infection results in robust macrophage and DC functions and enhanced Th_17_ polarization in the lung^27, 28^. Mice infected with a *hip1* mutant survived almost twice as long as wild-type (WT) Mtb-infected mice and had markedly reduced lung immunopathology, despite equivalent bacterial burdens at 2 to 4 weeks post-infection (wpi), and high levels of bacterial replication in the lungs of both groups of mice at 20 wpi. Therefore, the *hip1* mutant provided us with a unique opportunity to delineate the early lung immune environment that precedes the development of protective responses that are more favorable to the host than the pathogenic outcomes mediated by WT Mtb.

In this work, we identify mitochondrial metabolism and function in the early lung as critical for shaping subsequent host immunity to Mtb. Using single-cell RNA sequencing (scRNAseq), high-dimensional flow cytometry, metabolomics, and extensive bioinformatics analyses, we profiled WT Mtb- and *hip1* mutant-infected lungs at 2 wpi, when bacterial burdens in the two groups were equivalent. We found highly divergent immunometabolic programming between the two groups, particularly in AMs, IMs, monocytes, and T cell subsets. While WT-infected lung cells were largely glycolytic, the absence of Hip1 led to significant upregulation of both glycolysis and mitochondrial metabolic pathways such as fatty acid β-oxidation (FAO), amino acid oxidation (AAO), and oxidative phosphorylation (OXPHOS), which resulted in significantly higher metabolic energy output relative to WT Mtb. Further, we show that maintaining intact mitochondrial integrity and metabolic processes is critical for generating the energy needed to fuel macrophage functions and innate-T cell signaling networks that promote protective immunity. Our studies reveal that Mtb restricts mitochondrial metabolism and function in the early infected lung, in part through Hip1, which allows the pathogen to manipulate host immunometabolism to its benefit and drive pathogenesis. Additionally, we discovered a core mitochondrial immunometabolic signature associated with protective outcomes in mice, non-human primates (NHP), and humans. Overall, our studies suggest that immunometabolic programming in the lung that enhances mitochondrial OXPHOS and ATP output in the early stages of Mtb infection tips the balance away from pathogenic immune outcomes and towards more protective lung immune environments.

## Results

### Divergent lung transcriptional programs between mice infected with WT and *hip1* mutant Mtb at 2 weeks post-infection

To compare the early lung immune landscape following WT Mtb and *hip1* mutant infection, we infected C57Bl/6 mice *via* aerosol with a low dose (∼100 CFU) of WT Mtb H37Rv (n=5), *hip1* mutant (n=5) or uninfected controls (UI; n=5) and harvested lungs after 2 wpi for immunophenotyping by multiparameter flow cytometry (**Fig. 1a**) to assess multiple myeloid and lymphoid cell types and subsets (**Supplementary Fig. 1a**). At 2 weeks, bacterial burdens in the lungs of both infected groups were equivalent (**Fig. 1b**). We concatenated 50,000 CD45^+^ cells from each sample to assess the distribution of immune cell types by high-dimensional reduction using Uniform Manifold Approximation and Projection (UMAP) and FlowSOM (FlowJo LLC) for cell clustering using selected phenotypic markers (**Fig. 1c and Supplementary Fig. 1b)**. We identified alveolar macrophages (AM; SiglecF+ CD11c^+^), interstitial macrophages (IM; CD64^+^ CD11b^+^), monocyte derived macrophages (MDM; CD64^+^ CD11b^−^), dendritic cells (DC; CD11c^+^ MHCII^+^), monocytes (Ly6C^+^), neutrophils (Ly6G^+^), T cells (CD3^+^/CD4^+^/CD8^+^), B cells (CD19^+^), and NK cells (CD161^+^). Overall, the distribution and frequencies of immune cells between the two groups were not significantly different (**Fig. 1d and Supplementary Fig. 1c)**, except for higher frequencies of DC and NK cells and lower B cell frequencies in the *hip1* mutant group.

**Fig. 1.**
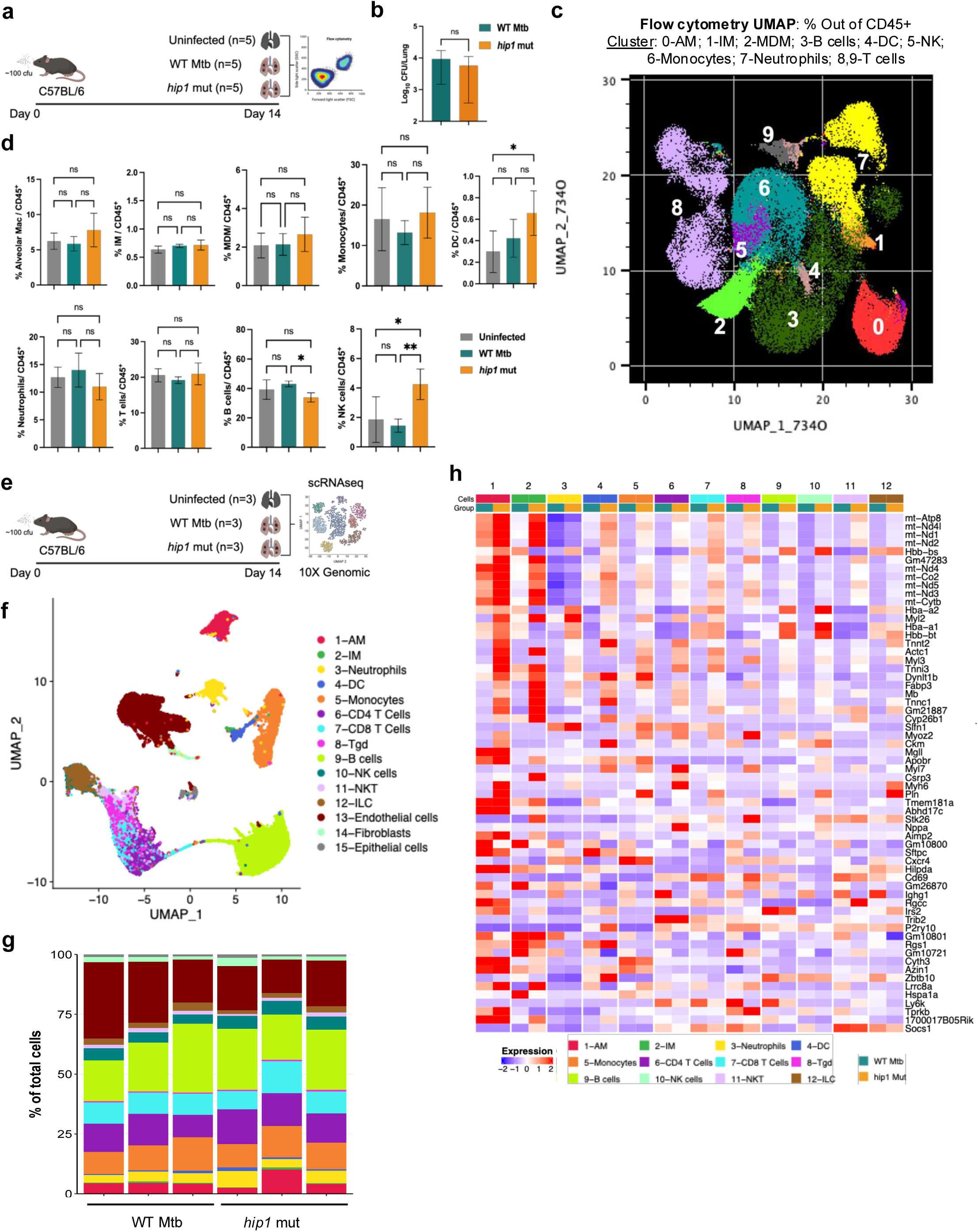
Immunophenotyping and single-cell transcriptomics of lung cells at 2 weeks post-infection in uninfected, WT Mtb- and *hip1* mutant- infected mice. **a,** Schematic overview of experimental design. Mice were left uninfected or infected with WT Mtb (H37Rv) and *hip1* mutant (n=5 per group) via aerosolization using ∼100cfu per animal. At week 2, lung cells were isolated and stained with a panel of antibodies to assess immune cell subsets by multiparameter flow cytometry. **b**, At 2 wpi, a section of the lung from each group was homogenized and plated on 7H10 agar plates. After 3 weeks of incubation, CFU from each sample was counted to determine lung bacterial burden in WT Mtb and *hip1* mutant groups. Data presented as mean ± SD and analyzed using paired t-test. Data are representative of 2 independent experiments. ^∗^p≤ 0.05; ns: p>0.05. **c**, Flow cytometry-based high-dimensional UMAP of concatenated CD45^+^ immune cells (50,000 events per animal) from uninfected, WT Mtb and *hip1* mutant groups (n=5/group). Cluster 0-9 was annotated based on cellular markers used in FLOWSOM map generation. **d**, Myeloid and lymphoid cell types identified by flow cytometry represented as the frequency of each cell type calculated out of the total concatenated CD45^+^ population (UMAP cluster) from the lungs of uninfected, WT Mtb and *hip1* mutant groups. Data are presented as mean ± SD and analyzed using a one-way ANOVA with Tukey’s multiple comparisons test. ^∗^p≤ 0.05; ^∗∗∗∗^ p≤ 0.0001; ns: p>0.05. **e**, Experimental design for aerosol infection and single-cell transcriptomics. Animals were infected by aerosolization of ∼100 cfu of either WT Mtb or *hip1* mutant or left uninfected (n=3 per group). At 2 wpi, lung cells were isolated, counted, and processed for scRNAseq (10X Genomics), droplet-based GEM bead capture, and library preparation, followed by single-cell sequencing and data analysis. **f**, scRNAseq-derived concatenated UMAP of Uninfected (n=3), WT Mtb (n=3), and *hip1* mutant (n=3) showing 15 different cell types in the mouse lung identified using reference-based annotation. **g**, Distribution of cell populations in the lungs of WT Mtb and *hip1* mutant. Each bar represents one animal. Samples are colored based on individual cell populations. **h**, Heatmap of differentially expressed genes in *hip1* mutant vs WT Mtb in the lung. Colors represent the Z-scores (row normalization) of the average expression of each gene for each group and cell type. Genes with high expression are shown in red and low expression in blue. Panel **a** created with BioRender.com.

We next sought to compare the transcriptional responses of lung immune cells at 2 wpi in WT-and *hip1* mutant-infected mice by conducting scRNAseq (10X Genomics). We infected C57BL/6 mice with a low dose (∼100 cfu) of aerosolized WT Mtb (n=3) or *hip1* mutant (n=3) and after 2 weeks, isolated single cell suspensions from the lungs of infected and uninfected (n=3) mice and subjected samples to droplet-based capture for 10X Genomics and subsequent single cell transcriptional profiling (**Fig. 1e**). Our data analysis included filtering out cells with < 200nCount, > 4500nCount, and >10% mitochondrial genes (**Supplementary Fig. 1d**). We captured up to ∼40,000 cells from nine animals to generate the global cellular map. Further analysis of the combined data resulted in 15 different cell types using the Seurat pipeline for generating a dimensional reduction UMAP (**Fig. 1f, Supplementary Fig. 1e** and **Supplementary Table 1**). Cell clustering was based on the mouse reference database^31^ as follows: AMs (*Ptprc, Mertk*, *SiglecF* and *Itgax)*, IMs (*Cx3cr1*, *Itgam)*, neutrophils ( *S100a8*), DCs (*Itgax, Ccl17*), monocytes (*Itgam, Csfr1*), CD4^+^ T cells ( *Cd3d*, *Cd4*), CD8^+^ T cells (*Cd3d*, *Cd8a*), gdT cells (*Cd3d*, *Trdc*), B cells (*Cd19*, *Cd79a*), NK cells (*Nkg7*, *Gzmb*), NKT cells (*Rora*, *Tbx21*, *Cd3d*), ILC (*Ncr1*), endothelial cells (*Pecam1*), fibroblasts (*Col3a1*, *Dcn*) and epithelial cells (*Sfptb*) (**Supplementary Fig. 1e, f**). While both infected groups showed similar distributions of cell populations (**Fig. 1g**), using cutoff criteria of fold change > 1.3 and < −1.3, and adjusted FDR p-value <0.05, we found several genes that were differentially expressed in the *hip1* mutant-infected lung compared to WT across multiple immune cell types, as shown in a heatmap (**Fig. 1h**). These data indicate significant divergence of transcriptional profiles between WT and *hip1* mutant groups as early as 2 wpi, when bacterial burdens are comparable.

### Upregulation of mitochondrial immunometabolic programs in *hip1* mutant-infected lungs

We performed differential gene expression analysis on the scRNAseq data to identify genes that were upregulated and downregulated in *hip1* mutant *versus* WT Mtb-infected lungs across all immune cell types. Using cutoff criteria of fold change > 1.3 and < −1.3, and adjusted FDR p-value <0.05, we found 39 significantly upregulated and 22 significantly downregulated genes in the *hip1* mutant-infected lungs compared to WT (**Fig. 2a**). Strikingly, the majority of upregulated genes (*mt-Atp8*, *mt-Atp6*, *mt-Nd2*, *mt-Nd3*, *mt-Nd4*, *mt-Nd4l*, and *mt-Nd)* corresponded to mitochondrial OXPHOS and the electron transport chain (ETC), metabolic pathways that are essential for ATP generation. This increase in genes related to mitochondrial bioenergetics and function was seen in multiple immune cells, including AMs, monocytes, neutrophils, CD4 and CD8 T cells, and NK cells (**Supplementary Fig. 2a-f)**. To capture a broader view of the pathways and biological processes in key immune cell types between the two groups, we conducted gene set enrichment analysis (GSEA) using the mouse molecular signature database (mSigdb) (**Fig. 2b**). Host metabolic pathways corresponding to cellular respiration, mitochondrial OXPHOS, ETC, and ATP synthesis were markedly enriched in the *hip1* mutant group. **Extended Fig. 1a,b** shows a visual representation of the OXPHOS pathway from the KEGG database globally and in AMs, with the mitochondrial proteins encoded by the upregulated genes depicted in red, including ND1, ND2, ND3, ND4, ND4L, ND5 (NADH dehydrogenases; complex I), cytochrome b (complex III), cytochrome c oxidases COX1-3, COX7A-B, COX8 (complex IV) and F-type ATPase-8 (complex V). Similar pathway representations were found for IMs, monocytes, T cells, and NK cells (**Supplementary Fig. g-j**). Together, these data suggest that Mtb fails to induce critical mitochondrial metabolic programming in the lung, in part through Hip1.

**Fig. 2.**
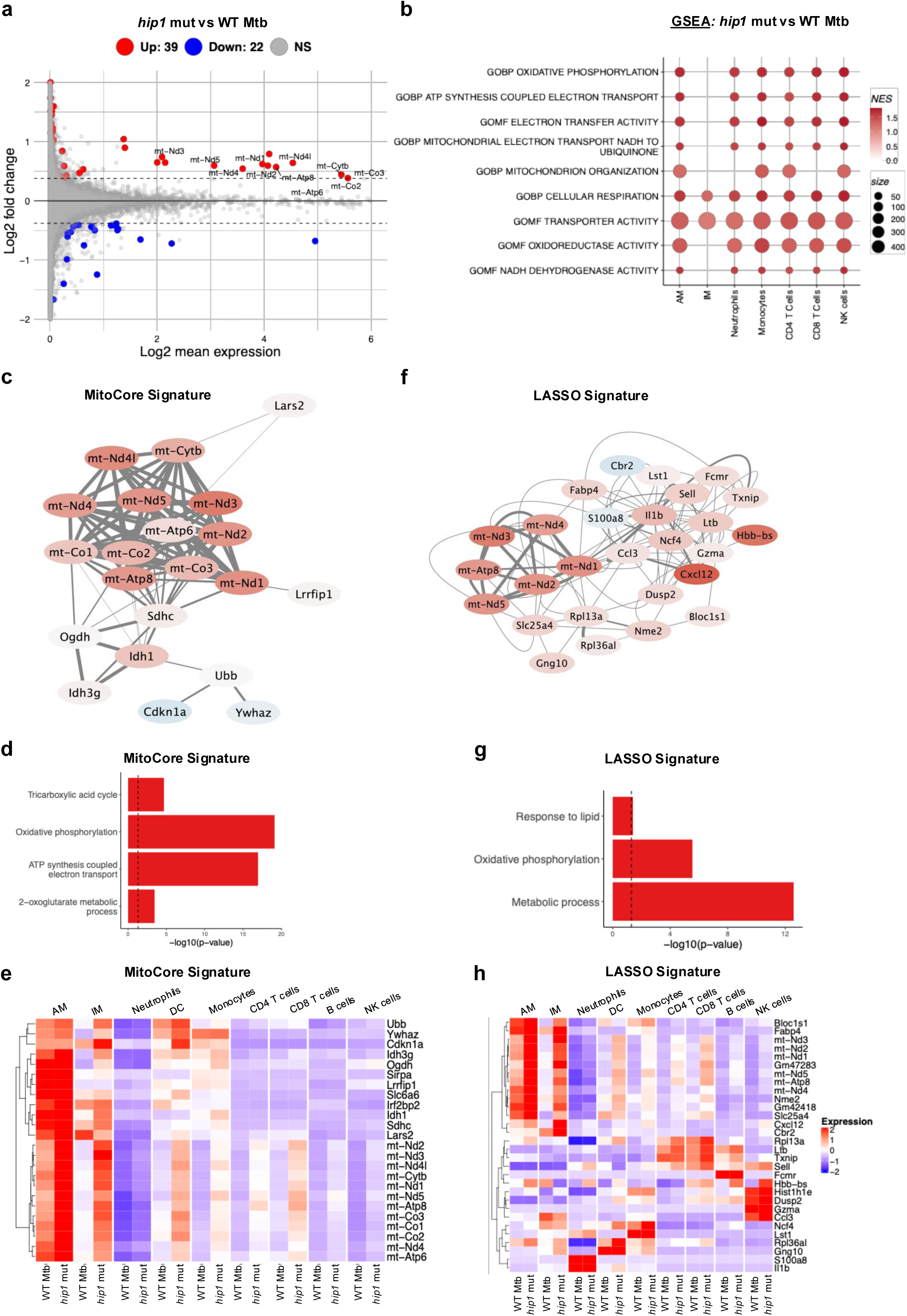
Differential gene expression analysis and derivation of core gene signatures using GCNA and LASSO across multiple cell types between *hip1* mutant- and WT Mtb-infected lungs. **a,** MA plot shows differential gene expression (DEG) profile in *hip1* mutant vs WT Mtb (n=3 per group). All cell populations except endothelial cells were included in DEG analysis. The x-axis shows the normalized log_2_ mean expression of each gene, and the y-axis shows the log_2_ fold change. Genes upregulated in the *hip1* mutant relative to WT are shown in red and downregulated genes are shown in blue. Genes with fold change greater than 1.3 or less than −1.3 and FDR adjusted p-value < 0.05 were considered to be differentially expressed. **b,** Bubble plot shows the pathways that were significantly enriched in *hip1* mutant-infected lungs compared to WT Mtb obtained from gene set enrichment analysis (GSEA). Pathways with p-value < 0.05 and NES > 0 were selected. A complete list of pathways is available in the supplementary data files. Pathways upregulated in *hip1* mutant vs WT Mtb are shown in red with intensity ranging from light red (low NES) to dark red (high NES). The size of the bubble represents the size of the reference pathway (i.e., the number of genes that comprise that pathway). **c**, Gene interaction network representing the core mitochondrial signature (MitoCore) derived from co-expression network analysis of scRNAseq data (n=3 per group). The lines represent evidence of connectivity between the genes as identified using StringDB, and the thickness of the lines represents the strength of the connection. The circles are colored based on fold change between the *hip1* mutant compared to WT Mtb, ranging from low (blue) to high (red). **d**, Pathway analysis of the MitoCore signature. The x-axis shows the -log_10_ p-value and pathways enriched in the signature are shown on the y-axis. The dotted line shows the 0.05 p-value cutoff. **e**, Heat map of genes in the MitoCore signature. Colors represent the Z-scores (row normalization) of the average expression of each gene for each group and cell type. Genes with high expression are shown in red and low expression in blue. **f**, Gene interaction network of the gene signature derived using LASSO analysis of scRNAseq data (n=3 per group). The lines represent evidence of connectivity between the genes as identified using StringDB, and the thickness of the line represents the strength of the connection. The circles are colored based on fold change between the *hip1* mutant compared to WT Mtb, ranging from low (blue) to high (red). **g**, Pathway analysis of the LASSO gene signature. The x-axis shows the -log_10_p-value and enriched pathways are shown on the y-axis. The dotted line shows the 0.05 p-value cutoff. **h**, Heat map of genes in the LASSO signature. Colors represent the Z-scores (row normalization) of the average expression of each gene for each group and cell type. Genes with high expression are shown in red and low expression in blue.

To complement and extend our analyses, we sought to identify core transcriptional signatures that would clearly distinguish WT- and *hip1* mutant-infected lungs using computational approaches such as gene co-expression network (GCN) and least absolute shrinkage and selection operator (LASSO) analyses. GCN analysis using CS-CORE identified 7 clusters (**Extended Fig. 1c**). Of these, cluster 2 strongly correlated with the *hip1* mutant (R^2^>0.5, p-value <0.05). String analysis showed this cluster to be enriched in mitochondrial genes *mt-Nd1*, *mt-Nd2*, *mt-Nd3*, *mt-Nd4*, *mt-Nd5*, *mt-Atp6,* and *mt-Atp8,* which we have termed the mitochondrial core signature (MitoCore, **Fig. 2c**). KEGG pathway enrichment of the MitoCore genes showed that the tricarboxylate (TCA) cycle, OXPHOS, and ATP synthesis pathways were significantly over-represented (p-value < 0.05) in the *hip1* mutant group compared to WT (**Fig. 2d**), particularly in AMs, IMs, DCs, monocytes, and T cells (**Fig. 2e**). In parallel, using the LASSO regression method, we identified 30 genes with positive coefficients that strongly distinguished the *hip1* mutant from WT Mtb-infected lungs. The string network showed high connectivity between mitochondrial genes within the LASSO signature (**Fig. 2f**), such as *mt-Nd1*, *mt-Nd2*, *mt-Nd3*, *mt-Nd4*, *mt-Nd5, mt-Atp8,* as well as *Bloc1s*, which is involved in maintaining mitochondrial respiration and integrity ^32, 33^. The LASSO signature also included genes involved in proinflammatory and antimicrobial responses, such as *Ccl3*, *Cxcl12*, *Il1b*, *Gzma*, *Ltb,* and *Lst1,* and was significantly enriched for OXPHOS, lipid response pathways, and other metabolic processes **(Fig. 2g**), particularly in AMs and IMs (**Fig. 2h**). Together, data from **Figures 1 and 2** support the idea that WT Mtb restricts mitochondrial metabolism in immune cells, particularly within macrophage subsets, and that strong proinflammatory responses are associated with OXPHOS upregulation in the early infected lung.

### Metabolomics of infected lungs identifies mitochondrial metabolic pathways restricted by Mtb

Since analysis of scRNAseq data indicated highly divergent mitochondrial metabolic programming in the presence or absence of Hip1, we sought to further delineate the underlying metabolic processes. First, we used Compass, a flux balance analysis algorithm that uses single-cell transcriptomics data to map cellular contexts and predict metabolic states ^34^. Compass analysis found a pronounced enrichment of multiple energy-generating metabolic processes that utilize sugars, amino acids, and fatty acids in the lungs of *hip1* mutant-infected mice compared to WT, including glycolysis, TCA cycle, FAO, and AAO, globally and in macrophages (**Fig. 3a-c and Extended Fig. 2a-c),** as well as in monocytes and T cells **(Supplementary Fig. 3a, b**). Mitochondria are the major sites for cellular energy production, wherein metabolites from glycolysis enter the TCA cycle in mitochondria to generate reducing equivalents, which donate electrons to the mitochondrial ETC and generate ATP through OXPHOS **(Fig. 3d**). Thus, Compass analyses suggest substantially diminished mitochondrial metabolism in key immune cells within WT Mtb-infected lungs at the very early stages. To experimentally validate predictions of divergent metabolic processes in the two groups, we next conducted untargeted metabolomics on 2-week lungs from uninfected, WT Mtb- and *hip1* mutant-infected mice (n=5 per group).

**Fig. 3.**
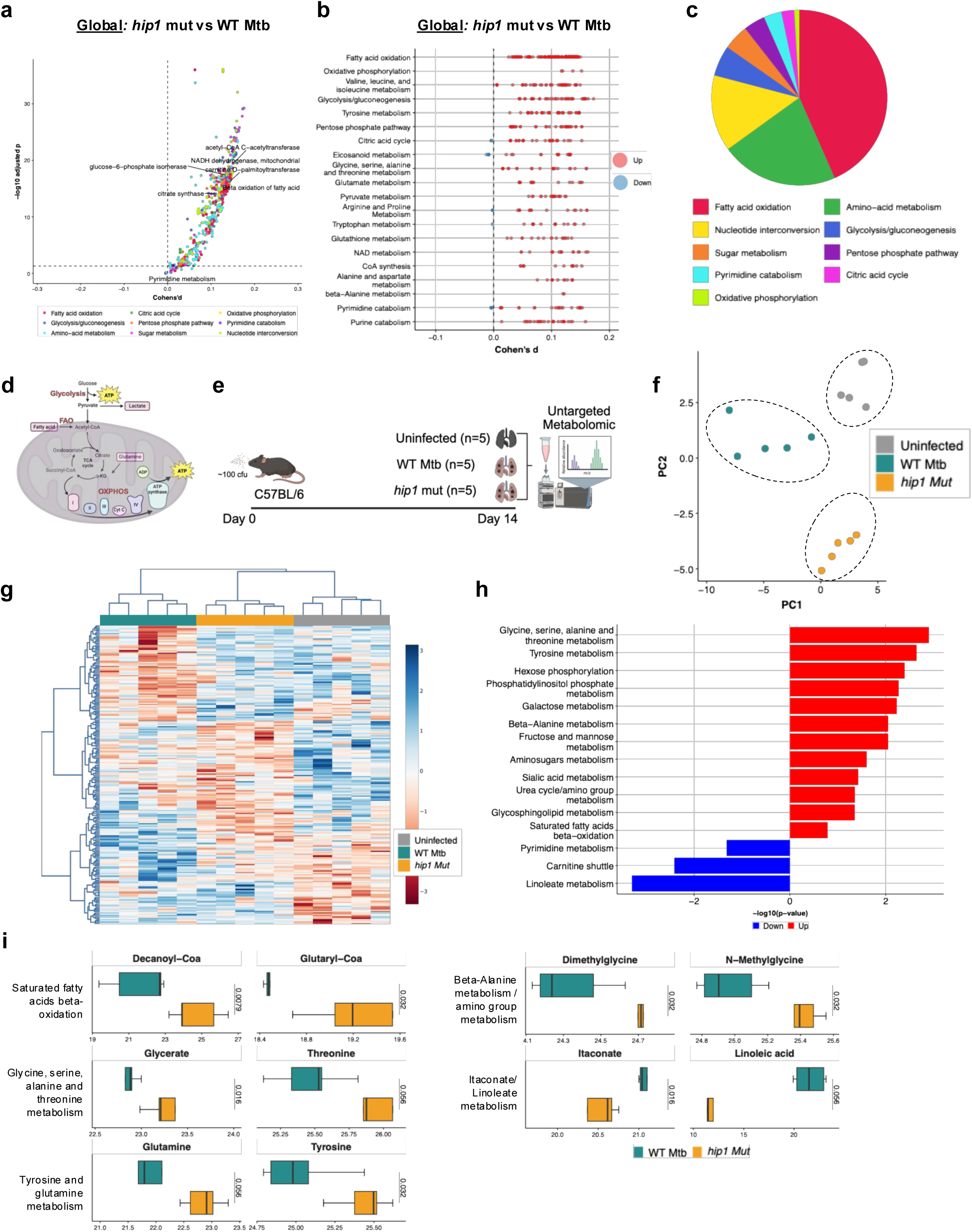
Metabolomic profiling of murine lungs at 2 weeks. **a**, Differential metabolic activity assessments between *hip1* mutant *versus* WT Mtb based on the Compass ^34^ score for each metabolic reaction, predicting multiple metabolic processes enriched in *hip1* mutant-infected lungs compared to WT Mtb. The effect size was assessed using Cohen’s d statistic, shown on the x-axis, calculated as the difference between sample means over the pooled sample standard deviation. Probability estimates are derived from Wilcoxon’s rank sum test, shown on the y-axis. Each dot represents one metabolic reaction colored by reaction subsystem (shown in c). **b**, Representation of metabolic reactions (listed on y axis) comparing the *hip1* mutant vs WT Mtb using Compass, where each reaction (dot) is partitioned by Recon2 pathway^108^ and colored by the sign of their Cohen’s d statistic value. Reactions with positive Cohen’s d (significantly enriched in *hip1* mutant) are shown in red, while negative Cohen’s d (enriched in WT Mtb) are shown in blue. **c**, Pie chart depicting the distribution of metabolic reactions grouped by Recon2 pathways, with Cohen’s d > 0 when comparing the *hip1* mutant *versus* WT Mtb. Individual reaction categories are labeled by the different colors shown. **d**, Graphical representation of mitochondrial metabolic pathways relevant to this study depicting glycolysis and OXPHOS interconnection via the TCA cycle (generated by Biorender). **e**, Experimental design for untargeted metabolomics studies from the lungs of uninfected, WT Mtb-infected, and *hip1* mutant-infected mice (n=5 per group) at 2 wpi. **f**, PCA plot of metabolomics data from uninfected, WT Mtb and *hip1* mutant groups. Significant genes with p-value < 0.05 calculated using ANOVA were used for PCA analysis. **g**, Heatmap showing hierarchical clustering of 228 significant metabolites (p-value < 0.05, ANOVA) in uninfected, WT Mtb, and *hip1* mutant groups. The colors represent the raw normalized Z-score, where metabolites present at high levels are shown in red and low levels are shown in blue. The hierarchical clustering of metabolite features was performed using Ward’s method^109^. **h**, Metabolic pathways upregulated (red) or downregulated (blue) in *hip1* mutant versus WT Mtb groups. X-axis shows p-value statistics obtained using the Mummichog pipeline^105^. **i**, Boxplot shows individual metabolites from the metabolomics data, representing key pathways enriched in *hip1* mutant *versus* WT Mtb-infected lungs. The following parameters are shown: minimum, lower quartile, median, upper quartile, and maximum. A two-sided Student’s t-test was used to compare sample means. Panel **d,e** created with BioRender.com.

Homogenized lungs were separated into cellular and extracellular fractions and each processed for metabolomics as described in Methods (**Fig. 3e**). The PCA plot and heat map show clear segregation of lung metabolomic profiles between the three groups, and hierarchical clustering of the metabolomics data showed distinct clusters of metabolites within each group (**Fig. 3f, g**). Categorizing the metabolites into pathways showed significant overlap with data from Compass analysis, with significant increases in catabolic metabolic processes in the lungs of *hip1* mutant-infected lungs, including AAO and FAO, amino sugar metabolism, and metabolism of hexose sugars such as glucose, galactose, fructose, and mannose (**Fig. 3h**). Notably, *hip1* mutant infection induced significantly higher levels of metabolites corresponding to mitochondrial metabolic processes such as AAO and FAO than WT Mtb, and lower levels of metabolites representing lipid transport and accumulation, such as the carnitine shuttle and linoleate metabolism (**Fig. 3h**). These data suggest that Mtb skews lung metabolism towards glycolytic metabolism and lipid synthesis while restricting breakdown of amino acids and fatty acids. **Fig. 3i** shows data for selected metabolites that were present at significantly higher levels in *hip1* mutant lungs, such as glycerate, threonine, glutamine, tyrosine, dimethylglycine, and N-methylglycine (AAO) and decanoyl-CoA and glutaryl-CoA (FAO), which fuel ATP generation through OXPHOS/ETC to provide the energy necessary for mounting robust immune defense against microbes^35^. In contrast, WT Mtb induced significantly higher levels of TCA cycle intermediates such as itaconate and linoleate, suggesting that the TCA cycle is disrupted^36, 37, 38^. Finally, integration of data from Compass analysis with our metabolomics data showed remarkable convergence between the metabolic processes identified by transcriptomics and metabolomics approaches (**Supplementary Fig. 3c**). Since robust mitochondrial metabolism is critical for mounting effective immunity against pathogens, our data reveal significant deficits in lung mitochondrial processes during WT Mtb infection, underscoring the critical role of mitochondria in driving immune outcomes.

### Distinct macrophage and monocytic populations drive mitochondrial metabolism at early stages of lung infection

Since macrophages are critical for shaping the early immune response to Mtb in the lung, we next sought to gain a deeper understanding of their immunometabolic profiles in the two infected groups. To hone in more precisely on the macrophage sub-populations across both groups, we filtered the scRNAseq data and identified 4 sub-populations of macrophages, annotated as AM1, AM2, AM3, and IM with distinct gene expression profiles (**Fig. 4a, b**). AM1 macrophages (which constituted the largest fraction at 60-65%) were enriched for lipid and FAO pathways and leukotriene metabolism, which generates pro-inflammatory mediators. AM2 macrophages (15-20%) had an activated M1-like proinflammatory profile and were enriched for antigen presentation and costimulatory functions, while the smaller (3-5%) proliferating (*Ki-67^+^*) AM3 subpopulation was enriched for cell cycle and nuclear division pathways. IMs (6-8%), which were defined based on expression of complement receptors and *C1qa* and *C1qb* (**Fig. 4c**), were enriched for processes associated with antigen-presentation, costimulation, cytokine production, and regulation of immune responses. While the frequency of each of these macrophage subpopulations did not differ between WT and *hip1* mutant groups (**Supplementary Fig. 4a**), GSEA pathway analysis revealed striking differences. The AM1, AM3, and IM subsets from the *hip1* mutant group were significantly enriched for pathways involved in mitochondrial biogenesis, OXPHOS, and cellular respiration (**Fig. 4d**), suggesting greater ATP output compared to WT. Additionally, IMs from the mutant group were significantly enriched for M1-like proinflammatory pathways such as IL-12 signaling, while also upregulating mitochondrial metabolic processes often associated with M2 macrophages, indicating a more balanced M1-M2 IM phenotype relative to WT. Pseudotime analysis, which infers the differentiation trajectories of cell subsets and their underlying gene expression programs, showed that AM2 and AM3 each arose from AM1, and that AM2 subsets likely differentiated further to generate IMs (**Fig. 4e**). Superimposing the MitoCore signature onto the AM1 to AM2 pseudotime trajectory showed that while AM1 subsets from the *hip1* mutant group acquired core mitochondrial programs needed to mount effector responses, AM1s from the WT group failed to express these gene programs (**Fig. 4f**). Similar results were seen for the AM1 to AM3 and AM1 to IM trajectories (**Fig. 4g, h**). Additionally, mitochondrial genes from the LASSO signature were also enriched in AM and IM subsets from the *hip1* mutant compared to WT **(Supplementary Fig. 4b-d)**. Taken together, we conclude that AMs and IMs induced by WT Mtb at early stages of lung infection are deficient in mitochondrial metabolic processes necessary for mounting effective and balanced immune responses against Mtb.

**Fig. 4.**
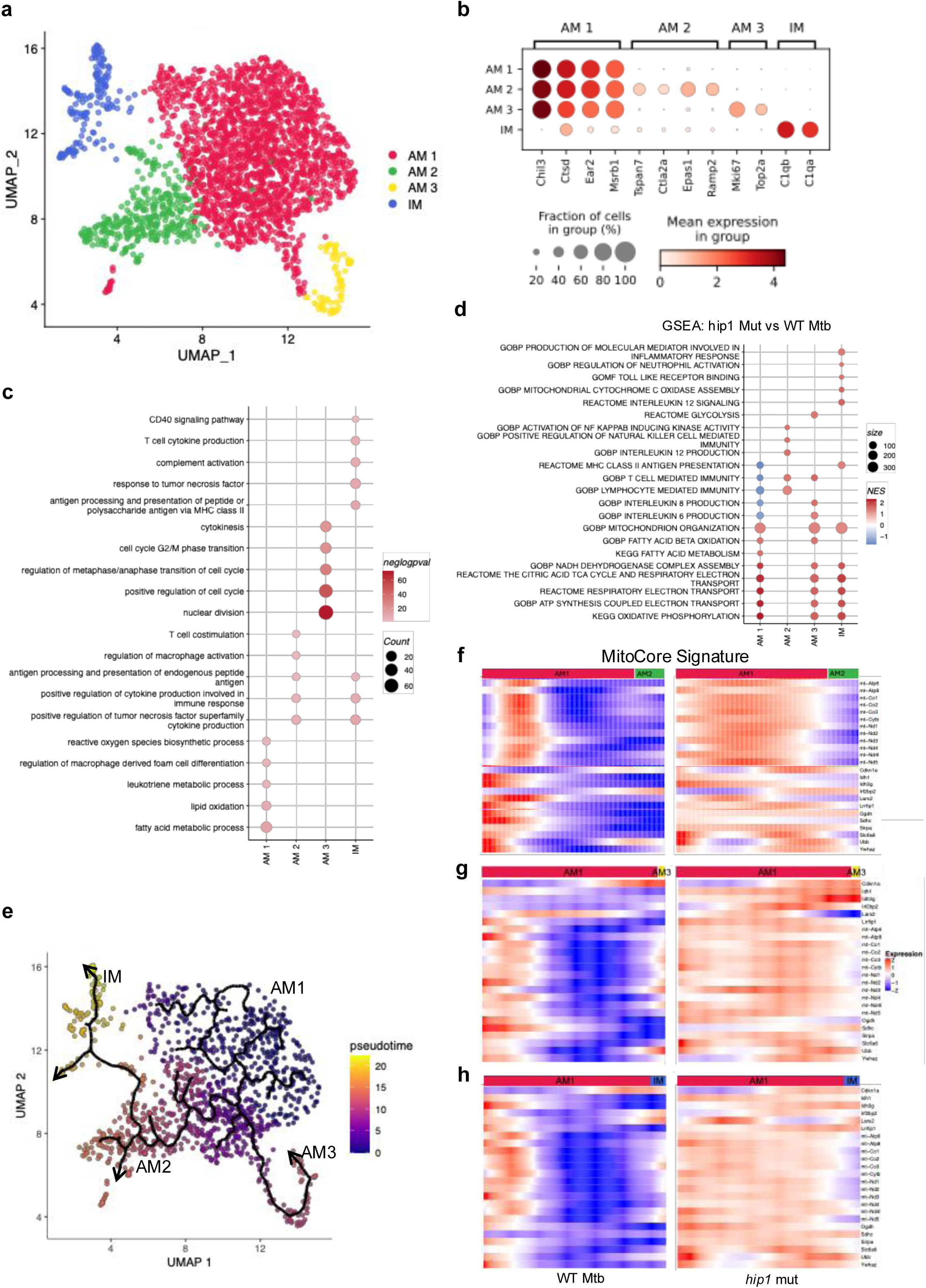
Phenotypic and functional signatures associated with macrophage subsets. **a**, UMAP projection of concatenated macrophages from all three groups (n=3 per group) segmented into alveolar macrophage (AM1, AM2, AM3) subsets and interstitial macrophages (IM) based on Seurat clustering using the nearest neighbor algorithm. **b**, Bubble dot plot shows expression of the key genes that classify each macrophage subset into AM1, AM2, AM3, and IM. The size of the dot indicates the fraction of cells expressing the gene, and the intensity of the color reflects the extent of expression of the gene. **c**, Bubble plot shows pathways differentially enriched in each macrophage subset compared to others. For each subset, genes with a fold change > 1.3 and FDR-adjusted p-value < 0.05 were selected for pathway analysis. Bubbles are colored based on p-value, where a darker color represents a lower p-value. The size of each bubble represents the number of genes in the reference pathway. **d**, GSEA showing pathways present in macrophage subsets from *hip1* mutant *versus* WT Mtb groups. Pathways with p-value < 0.05 obtained from GSEA analysis were selected for visualization. The bubble plot shows pathways upregulated (red) or downregulated (blue) in each subset from the *hip1* mutant compared to WT Mtb; the size of the bubble represents the number of genes in the reference pathway. **e**, Combined UMAP projection of *hip1* mutant and WT Mtb showing macrophage subsets in a pseudotime scale. The color represents the direction from early (violet) to later (yellow) timepoints in pseudotime, and arrows indicate the pseudotime trajectories. **f,g,h**, Heatmaps depicting the mitochondrial core gene signature in macrophage subsets on a pseudotime scale from **(f)** AM1 to AM2, **(g)** AM1 to AM3, and **(h)** AM1 to IM comparing *hip1* mutant vs WT Mtb. Each row represents a gene with the color gradient indicating expression levels (red for high, blue for low) of each in the indicated trajectory. The colored bar at the top indicates the specific macrophage subset annotation.

Monocyte-derived macrophages and other monocytic subsets that are recruited into the lung play an important role in the host antimicrobial response to Mtb. In all infected animals, we observed populations of monocytes with gene signatures that classified them into 3 distinct groups based on their differentiation states^39^, annotated as Mono1, Mono2, and Mono3 (**Extended Fig. 3a**). Ly6C^hi^ monocytes (Mono1) expressed high levels of *Ccr2* and *Ccl9,* which are known to enable infiltration into tissue and differentiate further into macrophages or DC (**Extended Fig. 3b**), showed enrichment in antigen presentation via MHC-II and in TNF-α signaling (**Extended Fig. 4c**). Ly6C^low^ monocytes (Mono2) expressed *Ace* and *Cd36*, reflecting pathogen uptake capacity (**Extended Fig. 3b)** and showed evidence of existing as resident monocytes in the vasculature with surveillance and myeloid/endothelial differentiation functions (**Extended Fig. 3c**). Pseudotime trajectory analysis showed that Mono1 subsets likely arose from Mono2 monocytes (**Extended Fig. 3e**). Finally, Mono3 subsets expressed granulocyte-associated genes such as S*100a8*, *S100a9,* and the neutrophil chemoattractant *Cxcl2* (**Extended Fig. 3b, c**). While monocyte subset frequencies were similar in both infected groups (**Supplementary Fig. 4e**), all three subsets from the *hip1* mutant were significantly enriched for mitochondrial biogenesis, ETC, OXPHOS, and ATP synthesis compared to WT Mtb (**Extended Fig. 3d**). Superimposition of MitoCore and LASSO signatures on monocyte pseudotime trajectories clearly showed higher expression of both signatures in monocytic subsets from *hip1* mutant-infected lungs (**Extended Fig. 3f and Supplementary Fig. 4f**). Overall, these data indicate that the altered mitochondrial metabolic programs seen in AMs and IMs extend to monocytic subsets in infected lungs.

### Impaired macrophage-T cell interaction networks in WT Mtb-infected lungs at 2 weeks

Antigen-specific IFN-γ^+^ CD4 T cell responses are typically detected ∼3 weeks after low-dose aerosol infection of mice^20, 40^. This delayed recruitment of T cells and their inability to fully control bacterial replication within macrophages is a result of Mtb’s immune evasion mechanisms that promote pathogenesis. Sub-optimal T cell functions are preceded by impaired innate immune activation and poor costimulation of T cells by antigen-presenting cells^15, 17, 25, 41^. We previously showed that the *hip1* mutant infection enhanced innate proinflammatory and costimulatory responses and augmented protective multifunctional Th_1_-Th_17_ responses in the lung compared to WT Mtb^24, 27, 28^. Bioinformatics analysis of T cell subsets and UMAP clustering identified CD4 and CD8 T cells corresponding to naive, activated, and memory subsets (**Fig. 5a and Supplementary Fig. 5a**) with comparable cell distributions in the two groups (**Supplementary Fig. 5b, c**). However, we found a striking enrichment of cytokine signaling pathways, including IL-1β, TNF-α, IL-12, IFN-γ, and IL-17 in activated CD4 T cells from *hip1* mutant-infected lungs (**Fig. 5b**), suggesting an earlier onset of protective Th_1_ and Th_17_ responses in the lungs of these mice. The significant under-representation of these pathways after WT Mtb infection suggests defective initiation of Th_1_ and Th_17_ responses and inadequate protective immunity. We next evaluated cell-cell communication networks between the different myeloid and T cell subsets using CellChat^42^, which employs curated databases of ligand-receptor pairs to score communication probabilities for various signaling pathways at a single cell level (**Fig. 5c**). CellChat analysis of 2-week scRNAseq data predicted strong interactions between AMs and monocytes and between AM/IM/monocyte subsets and CD4 T cells in pathways related to angiopoietin-like protein (ANGPTL) family members, and in prostaglandin (PGE2), leukotriene (LTB4), and ApoE signaling pathways in the WT group. Notably, these interaction networks were absent in lungs infected by the *hip1* mutant (**Fig. 5d-f**). Instead, the *hip1* mutant group had significantly higher relative information flow scores for signaling pathways associated with robust innate immune activation (MHC-II, TNF) and T cell costimulation (CD40). In addition, CellChat predicted strong interactions between AMs/IMs/monocytes and naïve CD4 T cells in the MHC-II signaling network and between CD40 on monocytes and CD40L on activated T cells, a costimulatory pathway that is known to be required for optimal Th_1_ and Th_17_ responses ^27, 28^. These networks were strikingly absent in WT at 2 wpi (**Fig. 5g-i**). Additionally, CD6 and BST2 signaling, which have been implicated in promoting Th_17_^43^ and DC activation^44^, respectively, were also absent in WT Mtb networks (**Fig. 5c**). Consistent with these data, we found earlier and significantly higher frequencies of ESAT6-specific IL-17^+^ T cells by ELISPOT, in the lungs of *hip1* mutant-infected mice starting as early as 3 wpi; differences that were even more prominent at 8 wpi (**Fig. 5j, k**). IFN-γ^+^ T cell frequencies were, however, comparable between the two groups. These results align with the known ability of Mtb Hip1 to dampen proinflammatory responses and limit Th_17_ polarization by impeding CD40-CD40L interactions^27, 28^. In summary, our analyses show that in the early infected lung, Mtb prevents crucial macrophage-T cell interaction networks that are essential for promoting development of protective T cell immunity and instead augments eicosanoid and lipid signaling pathways that have been linked to destructive inflammation and TB disease pathogenesis^45, 46^.

**Fig. 5.**
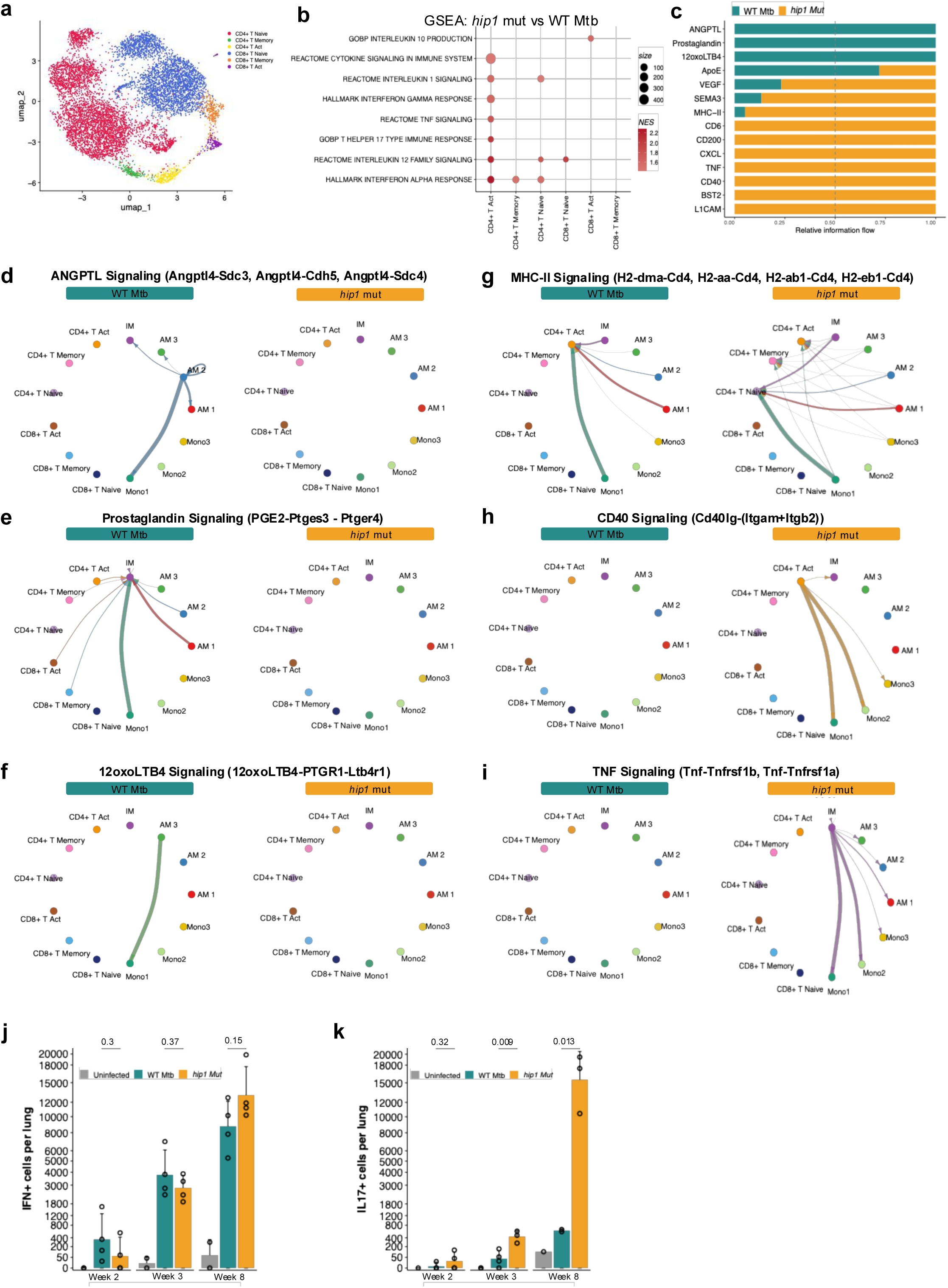
T cell subsets and cell-cell communication networks between myeloid and T cells. **a**, UMAP projection of concatenated T cells from the lungs of uninfected, WT Mtb- and *hip1* mutant-infected mice (n=3/group) segmented into different subsets based on Seurat clustering using the nearest neighbor algorithm. **b**, Gene set enrichment analysis (GSEA) showing pathways significantly enriched in T cell subsets from *hip1* mutant- *versus* WT Mtb-infected lungs. Pathways with p-value < 0.05 obtained from GSEA analysis were selected for visualization. The bubble plot shows pathways upregulated (red) or downregulated (blue) in the *hip1* mutant compared to WT Mtb for each subset; the size of the bubble represents the number of genes in the reference pathways. **c**, Cell-cell communication networks predicted by CellChat. The bar plot shows the strength of cell-cell interaction networks between myeloid and T cell subsets present in the *hip1* mutant (gold) vs WT Mtb (teal) group. The x-axis represents the relative flow of information calculated from the sum of cumulative probabilities between interacting cell types and scaled from 0 to 1. Higher values indicate stronger ligand-receptor interactions between the cells in each pathway; for e.g. stronger interactions that occur in *hip1* mutant-infected lungs (gold); TNF, CD40, and BST2 signaling networks are absent in WT Mtb (teal). Only pathways with communication probabilities <0.05 as obtained from CellChat analysis are shown. **d,e,f,g,h,i**, Representative cell-cell communication networks showing putative ligand-receptor pair interactions between macrophage, monocyte, and T cell subsets in *hip1* mutant- and WT Mtb-infected lungs. The arrow indicates the directionality of interaction, and the width of the line connecting the different cell subsets indicates the strength of communication; thicker lines indicate stronger interactions. **j,k**, ELISPOT assays quantifying the frequency of IFN-g^+^ **(j)** and IL-17^+^ **(k)** cells from mice infected with WT Mtb and *hip1* mutant (n=5 per group). Lung cells from weeks 2, 3, and 8 were plated in triplicate and stimulated with ESAT6_1-20_ peptide pools, followed by cytokine measurements as described in Methods. The difference in the means between the groups was assessed using a two-sided Wilcoxon non-parametric t-test.

### Mtb infection restricts OXPHOS and promotes mitochondrial dysfunction in alveolar macrophages

AMs are the first innate immune cells in the lung to encounter Mtb, and maintaining active mitochondrial metabolism and function is critical for generating sufficient ATP for mounting and regulating anti-TB host defense. To experimentally evaluate mitochondrial metabolism in AMs, we used Single Cell ENergetIc metabolism by profiIing Translation inHibition (SCENITH), which enables assessment of the metabolic state of individual cells in combination with flow cytometric-based immunophenotyping^47^. SCENITH evaluates the impact of metabolic inhibitors on cellular energetics by using protein synthesis (puromycin fluorescence) as a surrogate. Monitoring how protein translation changes when specific metabolic pathways are inhibited, and the resulting profile of translation inhibition, reveals the cell’s metabolic profile. We infected mice with either WT Mtb (n=8) or *hip1* mutant (n=8) for three weeks and treated isolated lung cells with 2-deoxy D-glucose (2DG), which inhibits glycolysis, or with oligomycin, which inhibits mitochondrial ATP synthase, for 30 minutes, followed by incubation with fluorescently-labeled puromycin for another 30 minutes, as described in Methods. We used multiparameter flow cytometry to phenotype AMs (**Supplementary Fig. 6a)** at the single cell level and measure puromycin mean fluorescence intensity (MFI) in control, 2DG- and oligomycin-treated lung cells from each group. AMs from the WT Mtb group showed a significant reduction in puromycin MFI following 2DG treatment but no significant change after oligomycin treatment (**Fig. 6a**, upper and lower plots), indicating that AMs from the WT Mtb group preferentially utilized glycolysis over OXPHOS. In contrast, AMs from the *hip1* mutant group showed a significant reduction in puromycin MFI following oligomycin treatment, but not with 2DG, compared to controls (**Fig. 6b**, upper and lower plots). This indicates that AMs from *hip1* mutant-infected lungs effectively utilized mitochondrial OXPHOS (**Fig. 6b**), which would lead to greater energy output than WT Mtb, as OXPHOS generates significantly more ATP than glycolysis. To investigate this further, we examined individual reaction fluxes predicted by Compass and evaluated the reaction scores for ATP synthesis in the two groups. Interestingly, we found significantly lower scores for ATP synthesis via complex V in WT Mtb for AM1 and AM3 subsets (**Fig. 6c**), but no differences in glucose uptake (**Supplementary Fig. 6b**). These data suggest that AMs from WT Mtb-infected lungs were defective in their capacity to utilize mitochondrial OXPHOS to generate ATP and were predominantly glycolytic, consistent with our SCENITH data (**Fig. 6a, b**). To assess the relative dependence on OXPHOS for ATP generation in the two groups, we infected bone marrow-derived macrophages (BMDM) for 24 hours (MOI=3) and measured intracellular ATP levels in the absence or presence of oligomycin. Oligomycin treatment led to a significant reduction in ATP from BMDMs infected with the *hip1* mutant, indicating that these cells actively used OXPHOS for generating ATP (**Fig. 6d**). In contrast, WT Mtb-infected BMDMs were unaffected by oligomycin, indicating that they did not use OXPHOS to generate ATP and preferentially relied on glycolysis for energy generation.

**Fig. 6.**
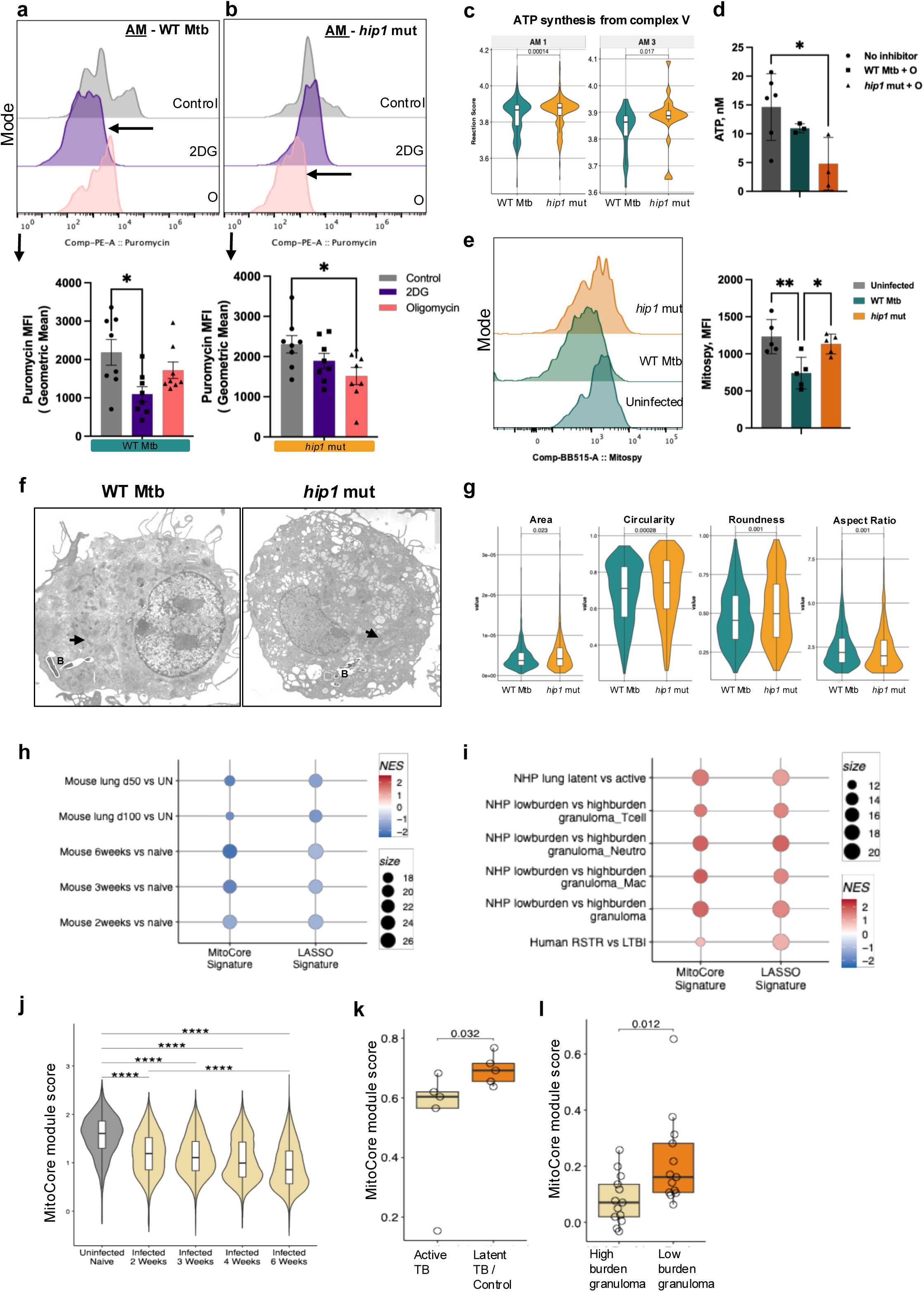
Single-cell profiling of macrophage metabolism and determination of mitochondrial morphometric features. **a, b**, Single cell metabolic profiling using SCENITH to analyze the metabolic state of lung alveolar macrophages from WT Mtb- **(a)** and *hip1* mutant-infected **(b)** mice at 3 wpi (n=8 mice/group). Flow cytometry of AMs after *ex vivo* treatment of lung cells with no inhibitor (control), 2DG, or oligomycin, showing puromycin MFI as histograms (upper) and geometric mean (below). Comparisons were made to the control uninhibited sample for each infected group. Data are presented as mean ± SD and were analyzed using one-way ANOVA with multiple comparison test. ^∗^p≤ 0.05; ^∗∗^p≤ 0.005; ns: p>0.05. Data are representative of two independent experiments. **c**, Violin and box plots of reaction scores obtained from Compass analysis for ATP synthesis from Complex V for AM1 and AM3 subsets. Scores were calculated for each cell from AM1 and AM3 subsets for WT Mtb and *hip1* mutant groups for the following parameters: minimum, lower quartile, median, upper quartile, and maximum. A two-sided Student’s t-test was used to compare the difference between sample means. ^∗^p≤ 0.05; ^∗∗^ p≤ 0.01; ^∗∗∗^ p≤ 0.001; ^∗∗∗∗^ p≤ 0.0001; ns: p>0.05. **d,** Concentration of intracellular ATP from BMDM infected with *hip1* mutant or WT Mtb treated with oligomycin (O) for 24 hours. The effect of oligomycin inhibition on ATP synthesis in the infected groups was compared against the control (pooled data from WT Mtb- and *hip1* mutant-infected macrophages with no inhibitor), measured in the cell lysate using the ATP ELISA kit (abcam 83355). Data are presented as mean ± SD and were analyzed using one-way ANOVA with multiple comparison test. (n=3/group) ^∗^p≤ 0.05; ^∗∗^p≤ 0.005; ns: p>0.05. Data are representative of two independent experiments. **e**, Flow cytometry histogram (left) and graphical representation of Mitospy-Green MFI to quantify mitochondrial mass of AMs from uninfected, WT Mtb- and *hip1* mutant-infected mice. (n=5 per group). Data are presented as mean ± SD and were analyzed using one-way ANOVA with Tukey’s multiple comparison test. ^∗^p≤ 0.05; ^∗∗^ p≤ 0.005; ns: p>0.05. The data are representative of two independent experiments. **f,** Electron microscopic analysis of BMDM infected with WT Mtb or *hip1* mutant for 24 hours. The arrow indicates the mitochondria, and “B” indicates intracellular bacteria. **g**, Mitochondrial morphology measurements obtained using ImageJ analysis of electron microscopy images from BMDMs following infection with either the *hip1* mutant or WT Mtb. Mitochondria were counted from 10-15 Mtb-infected cells per group using ImageJ/Fiji software. Violin and box plots show the following mitochondrial features: area, circularity, roundness, and aspect ratio. A box plot shows the following parameters: minimum, lower quartile, median, upper quartile, and maximum. The difference in the means between the *hip1* mutant and WT Mtb was estimated using a two-sided Wilcoxon non-parametric t-test. **h,i,j,k**, Mitochondrial metabolic signatures associated with protective outcomes in mice, NHP and humans**. (h)** Bubble plots show enrichment scores of the MitoCore and LASSO signatures in TB mouse lung scRNAseq datasets as described in **Supplementary Table 2**. The y-axis indicates the specific conditions being compared. The enrichment scores and corresponding p-values were calculated using GSEA. Higher scores are shown in red, and lower scores are shown in blue. Darker shade corresponds to a higher enrichment score.The size of the bubble reflects the size of the genes in the signature that mapped to the datasets. **(i)** Bubble plots show enrichment scores of the MitoCore and LASSO signatures in TB lung NHP and human datasets as described in **Supplementary Table 2**. The y-axis indicates the specific conditions being compared. The enrichment scores and corresponding p-values were calculated using GSEA. Higher scores are shown in red, and lower scores are shown in blue. Darker shade corresponds to a higher enrichment score. The size of the bubble reflects the size of the genes in the signature that mapped to the datasets. **(j)** Violin plots of the mitochondrial module score (calculated using AddModuleScore function from R package Seurat using the MitoCore signature genes as input) quantified in the mouse lung kinetics dataset comparing Mtb-infected lungs at week 2, 3, 4, and 6 with uninfected. A box plot shows the following parameters: minimum, lower quartile, median, upper quartile, and maximum. The difference in the means between the groups was estimated using a two-sided Wilcoxon non-parametric t-test. ****, p < 0.0001. **k, l,** Box plots of the mitochondrial module score in NHP datasets: Active TB vs LTBI (**k**) and high-burden vs low-burden granulomas **(l**). A box plot shows the following parameters: minimum, lower quartile, median, upper quartile, and maximum. The difference in the means between the groups was estimated using a two-sided Wilcoxon’s non-parametric t-test.

We next evaluated mitochondrial mass, which corresponds to the total amount of mitochondria in a cell and reflects mitochondrial biogenesis, degradation, and the cell’s capacity to generate ATP through OXPHOS^48, 49^. A higher mitochondrial mass generally indicates a greater capacity for ATP production. We stained lung cells from 3-week-infected mice (n=5 per group) with MitoSpy Green FM, which selectively labels mitochondria; measuring the accumulation of MitoSpy (by assessing MFI) provides a readout of mitochondrial mass. Interestingly, the MFI of MitoSpy in AMs (**Fig. 6e**, left and right plots) from WT-infected lungs was significantly lower than in uninfected mice, indicating that WT Mtb infection led to decreased mitochondrial metabolic capacity. However, mitochondrial mass in AMs from *hip1*-mutant-infected lungs did not decrease after infection, indicating preserved mitochondrial functions.

To further assess mitochondrial function and cellular health, we performed electron microscopy to examine mitochondrial ultrastructure and evaluate key morphological characteristics. Mitochondrial morphology refers to the shape and structure of mitochondria, which can vary depending on cellular energy demands and is a proxy for mitochondrial function and bioenergetic output. We infected BMDM for 24 hr (MOI=3), and processed cells for electron microscopy, followed by quantification of morphological parameters from the images as described in Methods. Compared to WT Mtb, mitochondria from *hip1* mutant-infected macrophages were significantly larger (as measured by area), showed greater circularity and roundness, and lower aspect ratios, reflecting substantially more ATP generation (**Fig. 6f, g**). Taken together, assessments of mitochondrial metabolic and functional states *in vivo* and *in vitro* provide strong experimental evidence of mitochondrial dysfunction in macrophages following Mtb infection.

### Mitochondrial metabolic signatures are associated with protective immune outcomes

We hypothesized that mitochondrial signatures preferentially expressed in the *hip1* mutant-infected lung could be associated with protective immune outcomes in different animal models and humans. We used the MitoCore and LASSO signatures to interrogate publicly available transcriptomics datasets from mouse, non-human primate (NHP), and human TB studies (**Supplementary Table 2**). To assess these signatures in mouse models of TB susceptibility and disease, we used data from Akter *et. al*., who studied the single-cell transcriptional profiles of WT Mtb-infected lungs in C57BL/6 mice at 50 and 100 days post-infection, and Pisu *et. al*., who published scRNAseq analyses of mouse lungs at 2, 3, 4, and 6 wpi. In both datasets^50, 51^, the MitoCore and LASSO signatures were significantly diminished in the lungs of Mtb-infected mice compared to naive mice (**Fig. 6h and Supplementary Fig. 6c**). To investigate whether MitoCore or LASSO signatures were enriched in datasets associated with host immune control and protection, we next analyzed single-cell transcriptomics data from two NHP studies. Esaulova *et. al.* compared lung cells from rhesus macaques with asymptomatic latent Mtb infection (LTBI) to those with symptomatic ATB disease^52^. Both the MitoCore and LASSO signatures were significantly enriched in the lungs of animals with LTBI compared to ATB (**Fig. 6i, and Supplementary Fig. 6d**). Gideon *et. al.* compared the transcriptomes of high-burden *versus* low-burden lung granulomas from cynomolgus macaques infected with Mtb, as proxies for immune environments that are permissive or restrictive, respectively, for bacterial growth^53^. Interestingly, we found highly significant enrichment of both signatures in low-burden granulomas that controlled bacterial growth, including within specific immune cell types such as macrophages, neutrophils, and T cells (**Fig. 6i, and Supplementary Fig. 6e**), showing that enhanced mitochondrial metabolism in the lung is associated with bacterial control and protective immune outcomes. This conclusion is further supported by our finding that the MitoCore signature was over-represented in individuals classified as “Resistors” (RSTR) compared to those with LTBI (**Fig. 6i**). RSTRs were defined as individuals with documented high exposure to TB who remained negative by IGRA (Interferon Gamma Release Assay), which measures Mtb-specific responses in blood^54^. **Fig. 6i** shows clear enrichment of MitoCore and LASSO signatures in monocytes from IGRA-negative RSTRs infected *ex vivo* with Mtb, compared to their counterparts from IGRA-positive asymptomatic individuals with LTBI, consistent with the prevailing hypothesis that RSTRs mount innate and/or adaptive immune responses that protect them from being infected with Mtb. Next, we aggregated the expression levels of the genes in the MitoCore signature to calculate a “module score” that quantifies the overall enrichment of each signature. Upon applying the MitoCore module score to the mouse datasets, we found that the module score in the lungs of infected mice decreased sequentially from 2 to 6 weeks as the mice progressed to TB disease (**Fig. 6j and Supplementary Fig. 6c**), indicating a negative association between the MitoCore module score and TB disease. Importantly, the MitoCore module score was significantly higher in NHP with low-burden granulomas and NHP with LTBI compared to those with high-burden granulomas or ATB disease (**Fig. 6k, l**), demonstrating a strong association between intact mitochondrial metabolism and protective immune outcomes against Mtb infection.

Overall, our studies reveal that impairing early mitochondrial metabolism and function is a key strategy employed by Mtb to promote an immunometabolic environment in the early lung, which enables the pathogen to prevent host protective immunity and favor pathogenesis. While the upregulation of aerobic glycolysis in macrophages by Mtb is well-established, our studies suggest that Mtb restricts OXPHOS and ATP synthesis and promotes mitochondrial dysfunction, leading to detrimental dependence on glycolysis in lung immune cells, which limits the generation of metabolic energy needed for mounting protective anti-TB immunity. Additionally, our finding that the early lung mitochondrial metabolic programs induced by the *hip1* mutant are linked to protective outcomes in humans and NHPs further underscores the critical need to maintain intact mitochondrial function in the lung to enable mounting protective immune responses against Mtb infection. Thus, targeting mitochondrial metabolic pathways for immunomodulatory therapies has the potential to ameliorate disease progression and lung damage. Moreover, vaccine strategies that enhance mitochondrial metabolism and function have the potential to induce efficacious immune signaling networks that better protect against TB.

## Discussion

Early events in the lung that shape protective immune responses to Mtb infection are not well understood but are critical for developing better vaccines and immunomodulatory therapies for TB^15, 55^. Here, we provide new insights that substantially shift our understanding of how virulent Mtb modulates early lung immunometabolism to its benefit. We show that maintaining intact mitochondrial metabolism and function in the early infected lung is pivotal for generating protective immunity and highlight an important role for Mtb Hip1 in driving pathogenic immune responses by restricting early OXPHOS and promoting mitochondrial dysfunction.

Cellular metabolism is a central regulator of the strength and duration of immune cell function in response to infection. Nutrients such as glucose, amino acids, and fatty acids directly impact the energy (ATP) generated by immune cells and shape their effector functions^56, 57^. Glycolysis rapidly generates energy and signaling molecules that promote proinflammatory and antimicrobial responses^58^. In addition, metabolic intermediates from glycolysis enter the TCA cycle in mitochondria to produce reducing equivalents that donate electrons to the ETC and generate ATP through OXPHOS via complex V^59^. Mitochondrial AAO and FAO provide further metabolic flexibility to fuel OXPHOS for ATP generation. Recent studies suggest that Mtb modulates host metabolism to its benefit, but we lack detailed knowledge of the early cellular mechanisms and immunometabolic networks that determine whether Mtb infection will result in detrimental immune responses and disease pathology or protective outcomes with minimal tissue damage^12, 60, 61, 62^. The prevailing model for Mtb regulation of host immunometabolism is that infection triggers metabolic reprogramming from OXPHOS to aerobic glycolysis, which promotes proinflammatory and antimicrobial macrophage responses that partially control Mtb during the early stages of infection. However, we know that sustained reliance on glycolysis for ATP generation, as seen in LPS-stimulated, M1-polarized macrophages, leads to disruption of the TCA cycle, suppression of OXPHOS, and inhibition of mitochondrial respiration^63, 64^. Conversely, M2-polarized macrophages, which promote anti-inflammatory responses and tissue repair, maintain an intact TCA cycle by relying on FAO and OXPHOS to meet their energy demands, thereby enabling efficient mitochondrial respiration. A major new insight provided by our studies is that Mtb’s reliance on glycolysis to generate ATP in the early infected lung is a consequence of mitochondrial dysfunction induced by Mtb, which leads to dysregulated lung inflammatory responses and tissue damage. In contrast, efficient use of mitochondrial metabolic processes such as AAO, FAO, and OXPHOS for ATP generation in the early infected lung, as seen in the absence of Hip1, leads to immune outcomes that are more protective. Since glycolysis plus OXPHOS provides 30-32 ATP per glucose molecule compared to a net output of 2 ATP from glycolysis alone, our studies support a model in which maintenance of intact mitochondrial metabolism and function is essential for providing the energy needed to induce a balanced M1/M2 response and innate-T cell interactions that promote protective immunity.

Another strength of our study is the experimental validation of immunometabolic pathways identified by bioinformatics analyses. Metabolomics studies confirmed striking enrichment of OXPHOS, AAO, and FAO metabolism in *hip1* mutant-infected lungs at 2 wpi. SCENITH analysis and ATP assays showed that AMs from *hip1* mutant-infected lungs actively utilized mitochondrial OXPHOS to generate ATP, while AMs from the WT Mtb group preferentially utilized glycolysis (**Fig. 6**). Moreover, reduced mitochondrial mass and aberrant morphologies of mitochondria from Mtb-infected macrophages provided further evidence for impaired mitochondrial health and ATP production. Mtb-infected lungs also harbored significantly higher levels of TCA cycle intermediates such as itaconate and linoleic acid, which reflect TCA cycle disruption (**Fig. 3**). Accumulation of TCA metabolites has been described in people with ATB^65, 66^, suggesting that the development of TB disease in humans is associated with aberrant mitochondrial metabolism. Interestingly, itaconate suppresses LPS-induced macrophage proinflammatory responses by inhibiting succinate dehydrogenase-mediated oxidation of succinate^67^. Since succinate controls IL-1β expression, HIF-1α activity, and ROS production^68^, the accumulation of itaconate in WT Mtb-infected lungs indicates early inhibition of proinflammatory responses. Notably, the LASSO signature that was associated with protective immunity was enriched for mitochondrial genes as well as genes that reflect robust innate responses, such as *IL-1b, Cxcl12,* and *Ccl3,* mitochondrial integrity (*Bloc1s1*), and *Fabp4*, which promotes FAO via transporting lipids into mitochondria (**Fig. 2**)^33, 69, 70, 71, 72, 73^. However, this signature was significantly underrepresented in AMs and monocytes from WT Mtb-infected lungs, consistent with our previous studies showing that Mtb dampens TLR2-dependent IL-1β, IL-6, and TNF-α production in macrophages through Hip1-mediated proteolysis of its substrate^29, 74, 75^.

We also found significantly higher levels of fatty acids such as decanoyl-CoA, glutaryl-CoA, and linoleic acid, and amino acids such as glutamine, tyrosine, and threonine in WT Mtb-infected lungs compared to the *hip1* mutant, indicating that catabolic metabolites that are vital for generating ATP via FAO, AAO, and OXPHOS were being actively utilized in mutant-infected lungs^35^. Since glycolytic M1 macrophages are capable of overcoming TCA cycle disruption by switching from glucose to glutamine metabolism, induction of glutaminolysis could be the mechanism by which mitochondrial functions are preserved in the absence of Hip1. However, the inability of Mtb-infected M1 macrophages to initiate glutaminolysis, combined with the high levels of fatty acids and linoleate metabolism in WT Mtb-infected lungs, likely leads to mitochondrial dysfunction, lipid synthesis, and immune suppression of T cell function^76, 77^. Indeed, linoleic acid is known to promote triacylglyceride (TAG) synthesis and lipid-rich foamy macrophages by functioning as the primary precursor for arachidonic acid, which cells convert into eicosanoids like prostaglandins (e.g., PGE2) and leukotrienes (e.g., LTB4), that typically have immunosuppressive and protective effects, respectively^78, 79^. In TB, PGE2 can be either immunosuppressive or protective depending on the stage of infection. Therefore, the timing of PGE2 and LTB4 production, and the ratio between the two, is critical for regulating the balance between pathogenic and protective immunity. Cell-cell communication analysis from WT Mtb-infected lungs showed strong interactions between IMs and AMs or Mono1 subsets in the PGE2 signaling pathway, and between AM3 and Mono1 subsets in the 12oxoLTB4 pathway (**Fig. 5**). 12oxoLTB4 is a key biologically inactive metabolic product of prostaglandin reductase (PTGR1)-mediated oxidation of LTB4, and its production suppresses the proinflammatory and chemoattractant functions of LTB4^80^. Neither the PGE2 nor 12oxoLTB4 pathways were present in the *hip1* mutant lungs at 2 wpi. Instead, *hip1* mutant-infected lungs harbored interactions between different macrophage and monocytic subsets in the TNF-α pathway, had greater interactions between macrophages and naïve CD4 T cell subsets in the MHC-II signaling pathway, and between activated CD4 T cells and monocytes in the CD40-CD40L costimulatory pathway (**Fig. 5**). The presence of these macrophage-T cell interaction networks denotes robust antigen presentation and costimulatory capacity, which are necessary for initiating effective Mtb-specific T cell activation and function^27, 28^. The absence of these critical immune networks in WT Mtb-infected lungs is consistent with the suboptimal and delayed T cell responses in Mtb-infected lungs. Thus, we propose that induction of early arachidonic acid metabolism and TAGs by WT Mtb leads to lipid-rich foamy macrophages within chronic lung lesions, an environment that is conducive to Mtb survival. Relevant to this idea, *hip1* mutant-infected lungs had significantly fewer foamy macrophages than WT Mtb with greatly reduced lung immunopathology^45, 46^, suggesting that metabolic rewiring towards OXPHOS and FAO in the early lung promotes tissue remodeling and repair at chronic stages. Interestingly, we found increased angiopoietin-like signaling (ANGPTL4) between AMs and monocytes in WT but not in *hip1* mutant groups (**Fig. 5**). ANGPTL4 is downstream of HIF-1α and, through its interactions with syndecans, prevents clearance of plasma triglycerides. Inhibiting ANGPTL4 is being explored as a therapeutic strategy for reducing triglyceride levels and improving cardiovascular outcomes^81, 82^. Since ANGPTL4 is induced via HIF-1α, which promotes glycolysis and suppresses mitochondrial respiration, we propose that inhibiting ANGPTL4 during TB may have favorable therapeutic outcomes.

The absence of CD40 costimulatory signaling networks in WT Mtb lungs at 2 wpi supports our previous publications, which showed that Hip1 impairs CD40-CD40L interactions and Notch signaling pathways. Further, CD40 agonism augmented Th_17_ polarization and enhanced protection, with IL-17 frequencies negatively correlating with Mtb lung burden^27, 28^. Here, we show that mice infected with the *hip1* mutant induce earlier and greater ESAT6-specific IL-17 responses in the lung over time (**Fig. 5**). Interestingly, a recent study showed that triggering CD40 signaling in LPS-induced M1 macrophages led to extensive metabolic reprogramming toward OXPHOS, FAO, and glutamine metabolism, pathways that are also preferentially induced by *hip1* mutant infection. CD40 agonism in tumor-associated macrophages induced glycolysis and proinflammatory responses without disruption of the TCA cycle and augmented anti-tumorigenic immunity^83^. Given our findings that the *hip1* mutant showed a similar profile, i.e., upregulation of proinflammatory responses without TCA disruption or impairment of mitochondrial metabolism, it is interesting to speculate that induction of CD40 signaling rewires lung immunometabolism toward a more beneficial M1/M2 macrophage phenotype. Assessing how CD40 engagement *in vivo* impacts lung metabolic programming and disease pathology in the context of WT Mtb infection will provide further mechanistic insights and is a topic of future studies for our group. Moreover, since work from several groups has highlighted the importance of inducing Th_17_ cells for vaccine-induced protection against TB, augmenting CD40 signaling and mitochondrial metabolism through adjuvants and small molecules, respectively, has the potential to improve vaccine efficacy.

We identified three transcriptionally distinct AM subsets and one IM subset in 2-week-infected lungs (**Fig. 3**), which were annotated based on the study by Pisu *et al*.^19^, who found that IMs were predominantly M1-like controllers while AMs were overall more M2-like and permissive for Mtb growth. However, both IM and AMs contained controller and permissive subpopulations. Although their study used high-dose infection with Mtb Erdman and sorted infected macrophages for scRNAseq, while we conducted scRNAseq on the whole lung after low-dose H37Rv infection, we observed significant congruence in the immune signatures from AM and IM subpopulations across the two studies (**Fig. 4**). At 2 wpi in our study, IMs from the *hip1* mutant-infected lungs were significantly enriched in pathways that promote innate immune activation, antigen presentation and inflammation, mitochondrial organization, and OXPHOS/ETC compared to WT Mtb, suggesting that higher energy generation from mitochondrial metabolism underlies the enhanced IM effector phenotype and functions (**Fig. 4**). Further, we identified AM subpopulations with immune signatures that strongly overlapped with both the controller and permissive phenotypes described in Pisu *et al*. Interestingly, the AM1 subset in our study was predominantly M2-like and exhibited a permissive phenotype in the WT group. However, *hip1* mutant infection led to a more complex pro- and anti-inflammatory M1/M2 signature that was previously shown to be associated with a controller phenotype^12, 19^, and was enriched for both MitoCore and LASSO signatures (**Fig. 4**). Thus, our study uncovers an important link between mitochondrial metabolism and beneficial immunity in the Mtb-infected lung and reveals that the early metabolic environment of the lung is a critical determinant of whether the immune response skews towards pathogenic or protective outcomes. Our data also provide new *in vivo* insights into previous reports of Mtb infection impacting mitochondrial metabolism. Using Seahorse assays, *in vitro* studies in human monocyte-derived macrophages and CD8 T cells showed decelerated mitochondrial bioenergetics and decreased rates of ATP production following Mtb infection^60, 84^. Further, metabolic profiling of lung cells from Mtb-infected mice at 3 wpi (high-dose intranasal Mtb Erdman infection) showed that AMs subsets were more dependent on glycolysis than mitochondrial metabolism^12^.

How might Hip1 contribute to restricting mitochondrial metabolism? We have previously demonstrated that Hip1 is a serine protease that dampens host immunity through proteolytic cleavage of its substrate GroEL2. Full-length GroEL2 protein was highly immunostimulatory to macrophages and DCs and augmented Th_17_ polarization, but cleaved GroEL2, the dominant form in Mtb, was poorly stimulatory in macrophages and DCs^30, 74, 85^. Interestingly, GroEL2 was shown to traffic into mitochondria within macrophages and interact with the mitochondrial protein mortalin, although the authors did not distinguish between full-length and cleaved GroEL2 in this study^86^. We speculate that Hip1-mediated cleavage of GroEL2 results in targeting of the cleaved form of GroEL2 to mitochondria, where it might function to impair mitochondrial functions through a yet undiscovered mechanism. Our finding that mitochondrial mass and morphologies were aberrant in WT Mtb infection supports the idea that cleaved GroEL2 drives mitochondrial dysfunction, while the absence of cleaved GroEL2 in the *hip1* mutant preserves mitochondrial function (**Fig. 6**). Additional Mtb proteins that have been detected in mitochondria include Rv1813^87^ and Rv0674^88^, but the underlying mechanisms are not known. Future studies warrant a comprehensive identification of Mtb effector proteins that impair mitochondrial metabolism and function. Additional Mtb mutants that have provided insight into how Mtb manipulates early lung immune responses include Δ*cpsA,* which showed delayed induction of T cell immunity and impaired recruitment and dissemination of bacteria from AMs to monocyte-derived cells^89^. CpsA was reported to inhibit NADPH oxidase (NOX) assembly, which mediates host defense through ROS generation but can also cause oxidative stress. Whether or not CpsA is targeted to NOX4 in mitochondria and/or modulates mitochondrial metabolism is, however, not known. The Δ*RD-1* mutant, which lacks the ESX-1 locus, was shown to be defective in exiting AMs and had impaired IL-1R and inflammasome signaling^90^. IL-1β is negatively regulated by NADPH oxidase, which is critical for mitochondrial bioenergetics^69, 72^, suggesting links between metabolism and immunity that remain to be explored. Although IL-1β is important for anti-mycobacterial immune responses, we know that Mtb limits inflammasome activation and IL-1β secretion through Hip1, Zmp-1, and PknF, as avirulent mutants that lack each of these factors induced much stronger inflammasome activation and IL-1β production^29, 91, 92^ relative to WT Mtb. Together, these data support a model in which restriction of mitochondrial metabolism by Hip1 and other virulence factors may be a common mechanism underlying the ability of Mtb to dampen IL-1β and other proinflammatory responses in the early lung and highlight that maintenance of mitochondrial function is beneficial to the host.

To investigate the relevance of our findings to the broader TB field, we applied the MitoCore and LASSO signatures to publicly available transcriptomic datasets from animal^50, 51, 52, 53^ and human studies^54^. In mice infected with WT Mtb, both the MitoCore and LASSO signatures were negatively associated with lung bacterial burden. Moreover, the MitoCore module score was significantly lower at 6 wpi compared to 2 and 3 wpi, suggesting that mitochondrial metabolic processes in the lung continue to deteriorate as disease progresses (**Fig. 6**). In NHP and humans, infection outcomes that result in asymptomatic LTBI and/or the RSTR phenotype are considered to reflect host immune control and protection against TB disease. Our finding that the MitoCore and LASSO signatures were highly enriched in NHP with LTBI compared to ATB and that the MitoCore module score was significantly higher in NHP granulomas with low bacterial burdens compared to high-burden granulomas (**Fig. 6**) strongly suggests that these animals and individuals retained robust mitochondrial metabolic activity in the early stages of Mtb infection, which in turn provided the necessary energy to mount protective immunity. Additionally, Mtb-infected monocytes from RSTRs were also significantly enriched for the MitoCore and LASSO signatures compared to individuals with LTBI^54^ (**Fig. 6**). Recently, the RSTR phenotype was shown to be associated with polymorphisms in the AMP-activated protein kinase (AMPK), a master metabolic regulator that boosts ATP production in response to stress via enhanced FAO and glucose uptake^93^ while decreasing fatty acid synthesis^94, 95^. This is strikingly similar to the lungs after *hip1* mutant infection at 2 wpi, which showed strong induction of FAO and AAO, downregulation of lipid synthesis pathways (**Fig. 3**), and enhanced macrophage-T cell interaction networks associated with protective immunity (**Fig. 5**). Thus, our studies in mice contribute new insights into immunometabolic pathways that drive immune control or clearance in NHP and humans, which merit further investigation. Mice are, however, not the ideal model to study asymptomatic infection and immune control. Both cynomolgus and rhesus macaque models have demonstrated that after low-dose Mtb infection, some of the infected animals remain asymptomatic while others progress to ATB. While it is not possible to predict at the early stage which animals will manifest as controllers or non-controllers at 3 to 4 wpi, assessing the MitoCore signature and metabolically profiling serial BAL and blood samples (via SCENITH or metabolomics), and correlating these data with clinical, microbiological, and pathological outcomes has the potential to yield important information.

We acknowledge that our study has limitations in that we were not able to differentiate between Mtb-infected and bystander cell populations in the lung. Mouse studies that successfully used fluorescently labeled Mtb or various Mtb reporter strains used high-dose infection in order to have sufficient numbers of infected macrophages to perform transcriptomics^12, 19, 62^ on flow-sorted macrophages. Here, we sought to capture the entire spectrum of immune cells and their interaction networks in the lung at a very early stage with a physiologically relevant low-dose model, which is not conducive to such assessments. We were also unable to directly measure cellular bioenergetics in lung cells using extracellular flux assays due to the unavailability of a Seahorse instrument in our BSL3 facility. As an alternative, we measured puromycin intensity after inhibition of glycolysis or mitochondrial OXPHOS, which has been shown to serve as a proxy for ATP synthesis^47^ (**Fig. 6**).

Overall, our studies reveal that the balance between protective and pathogenic immunity to TB is determined by mitochondrial metabolism in the early lung. Importantly, Mtb limits ATP generation by restricting mitochondrial FAO, AAO, and OXPHOS and induces mitochondrial dysfunction, which enables the pathogen to skew AM responses towards ineffective immunity and fatty acid synthesis, leading to lung immunopathology. Thus, targeting mitochondrial metabolic pathways has the potential to elicit beneficial immune responses through vaccination and/or reduce pathogenic immune responses and lung damage during ATB via adjunctive host-directed therapies.

## Acknowledgements

This work was supported by NIH R01AI134244 and R01AI155023-01A1 to J.R. and supported by NIH P30AI168386 and P51OD011132. We kindly acknowledge all former and current lab members of the Rengarajan lab and Day lab for their valuable input and support. We thank Kathryn Pellegrini, Gregory Tharp, and Steven Bosinger from the Emory Primate Center’s Genomics Core for 10X Genomics procedures and NextGen sequencing (supported in part by NIH P51 OD011132, NIH S10 OD026799), and the Robert P. Apkarian Integrated Electron Microscopy Core Facility (RRID: SCR_023537). Model figures were generated using BioRender (https://biorender.com).

## Author contributions

**Conceptualization**, H.K.D., P.B., J.R.; **Methodology**, H.K.D., P.B., A.B.E., L.B.H., R.C.G-F., J.C., J.R.; **Investigation**, H.K.D., P.B., L.B.H., S.D., A.P., T.R., J.K.S., R.M.L., S.L.G., T.J.W., J.C., J.R.; **Validation**, H.K.D., P.B., R.C.G-F., J.C., J.R.; **Data analysis and visualization**, P.B., H.K.D., M.C.K., A.S., A.P., L.B.H., J.K.S., K.E.P., J.C., J.R.; **Resources**, J.R.; **Writing – original paper**, H.K.D., P.B., J.R.; **Writing - Review & Editing**, H.K.D., P.B., L.B.H., S.D., A.P., T.R., J.C., J.R.; **Supervision** – J.R., **Funding acquisition**, J.R.

## Methods

### Mice

C57BL/6 wild-type female mice were purchased from The Jackson Laboratory. All mice used for experiments were eight to twelve weeks of age. Mice were housed in either the Emory Primate Center (EPC) animal ABSL-1 or ABSL-3 vivarium under sterile conditions with sterile food and water provided *ad libitum*. All animals were handled according to the regulations formulated by the Emory University Institutional Animal Care and Use Committee (IACUC).

### Bacterial strains

*Mycobacterium tuberculosis* (Mtb) strains H37Rv and H37Rv *hip1* mutant^75^ were used. As previously described^27, 28, 30, 74^, Mtb strains were cultured in liquid Middlebrook 7H9 (BD Difco) media supplemented with 0.5% glycerol (Sigma), 10% oleic acid-albumin-dextrose-catalase (OADC) (BD), and 0.05% Tween 80 (VWR) at 37 °C and shaking at 75 rpm. Additionally, 20 μg/mL kanamycin (Sigma) was included for growing the *hip1*mutant. Stocks were prepared by growing cultures to an OD_600_ of 0.4–0.6, then filtered and resuspended in 7H9 media with 25% glycerol (Sigma) and stored at −80C. Before using, stocks were titered to determine CFU.

### Aerosol infection of mice

Mtb cultures for aerosol infection were prepared as previously described^27, 28^. Briefly, mice were secured in separate glass canisters and infected via the aerosol route (∼ 100 CFU) using a nose-only exposure chamber (In-Tox Products). On one day following aerosol infection, 3 mice per group were euthanized, and lungs were harvested to determine bacterial burdens for each animal, and the rest were processed for downstream flow cytometry, transcriptomics, or metabolomics. For enumeration of bacteria, a portion of the lungs was harvested in sterile 2 mL tubes (Sarstedt) containing stainless steel beads (Next Advance) and PBS + 0.02% Tween80. The lungs were then homogenized in a Bullet Blender (Next Advance). Serial dilutions were plated onto Middlebrook 7H10 plates (with or without 20 μg/mL kanamycin for *hip1*mutant) to determine the CFU after 3 weeks of incubation at 37 °C.

### Mouse tissue harvest and isolation of lung cells

After euthanasia, each mouse lung was dissected carefully and transferred to a C-Tube (MiltenyiBiotec) containing 5 ml HBSS (Corning) supplemented with 2% heat-inactivated FBS (Gemini) and 10mM HEPES (Corning). A mixture of collagenase type IV (1mg/ml; Worthington) and DNAse I (10ng/ml; Worthington) was added to each tube. Tissue homogenization was performed inside the biosafety cabinet using an automated gentleMACS Dissociator (MiltenyiBiotec) using the manufacturer’s murine lung processing program and placed in a 37°C (with 5% CO_2_) incubator for 30 min. Following this time, the lungs were dissociated again using a murine lung processing program. After the run, tubes were detached from the dissociator, and the samples were centrifuged at 300*g* for 5 min at 4°C, after which the supernatant was removed, filtered or stored and used for other assays. Cell pellets were resuspended in 3 ml of RBC lysis buffer (Sigma) and incubated for 3 mins at room temperature, followed by washing with 15 ml of Incomplete RPMI media containing 10% FBS. The cell suspension was centrifuged at 1500 rpm for 5 min at 4°C and the supernatant was collected and stored at −20°C. Cell pellets were resuspended in 5 ml of Incomplete RPMI media, then allowed to pass through a 70-μm strainer. Cells were counted in Countess slides (VWR) using trypan blue dye exclusion. Subsequently, the cell suspension was centrifuged at 1500 rpm for 5 min at 4°C. The final cell pellets were resuspended in complete media for flow cytometry staining, 10X Genomics single-cell RNA sequencing, or other assays.

### Flow cytometry

For immunophenotyping of lung cells, cells were initially incubated with live/dead dye Fixable Near-IR Dead Cell Stain Kit (Invitrogen). Additionally, mouse Fc block (BD) was used before staining. The live/dead stain (1:1000) and Fc block (1:50) mixture was diluted in PBS (Sigma) and 100 μL per sample was incubated for 30 mins at 4°C. After 30 mins cells were washed in FACS buffer containing FBS and were stained with fluorescently-labeled antibodies for flow cytometry as follows: BUV496 Rat anti-Mouse MHC-II (clone: 2G9, BD), BUV563 Hamster anti-Mouse CD45 (clone: I3/2.3, BD), BUV661 Rat anti-Mouse CD8a (clone: 53-6.7, BD), BUV805 Rat anti-Mouse CD4 (clone: RM4-4, BD), BV480 Hamster anti-Mouse CD103 (clone: 2E7, BD), BV605 Rat anti-mouse Siglec-F (clone: E50-2440, BV421 Rat anti-Mouse CD64 (clone: X54-5/7.1, Biolegend), BV570 Mouse anti-mouse CD161 (clone: PK136, Biolegend), BV711 Hamster anti-mouse CD11c (clone: N418, Biolegend), BV750 Rat anti-mouse CD19 (clone: 6D5, Biolegend), FITC Rat anti-mouse Ly-6G (clone: 1A8, Biolegend), PE/ Dazzle 594 Rat anti-mouse Ly-6C (clone: HK1.4, Biolegend), PE/Cy5 Hamster anti-mouse CD3ε (clone: 145-2C11, Biolegend), PE-Cy7 Rat anti-mouse/human CD11b (clone: M1/70, Biolegend). The surface stain antibodies were diluted in FACS buffer (PBS [Sigma], 2% heat-inactivated FBS [Gemini], and 2 mM EDTA [Corning]). Following staining, cells were fixed in 2% paraformaldehyde (PFA, diluted in FACS buffer) (Electron Microscopy Sciences) for BSL-3 samples and placed in 4°C until acquisition (up to 24 hr post-staining). For compensation, Anti-Rat and Anti-Hamster Ig k/Negative Control Compensation Particles (BD), ArC Amine Reactive Compensation Bead Kit (Invitrogen) were used. All samples were acquired by flow cytometer BD A5 Symphony using FACSDiva (BD) software. All Data were analyzed using FlowJo software (FlowJo LLC).

### Flow cytometric high dimensional data analysis

Flow cytometric analysis was performed using FlowJo software (FlowJo LLC). First, data quality was checked for good events and filtered using the PeacoQC plugin. Second, compensation was applied using proper compensation controls, and then a downstream gating strategy was applied based on the panels designed. For high-dimensional reduction and clustering, FlowJo plugins were used to generate UMAP projections and FlowSOM clustering maps. In brief, an equal number of selected cell events were downsampled from each sample and then concatenated into one file. Keywords were included in each sample before concatenation for downstream identification. UMAP plugin generated different sizes of clusters based on the marker features used in the panel. To annotate clusters, we used the FlowSOM-generated color-coded clusters and heatmaps corresponding to each population. The comparison between each sample and group was made by quantifying the cell number or frequency in each cluster population.

### 10X Genomics scRNAseq GEM chromium capture and data analysis

After two weeks of infection, mice were euthanized, and the lungs were harvested and homogenized to obtain single cells. Cells were passed through a 70-μm strainer and resuspended in incomplete RPMI media at a concentration of approximately 1 million cells per ml. Cell counts were performed using Countess slides with viability assessed via trypan blue dye exclusion before GEM capture. Single cell suspensions were prepared and loaded onto the 10X Genomics Chromium Controller in the BSL3 facility using the Chromium NextGEM Single Cell 5’ Library & Gel Bead Kit v2 to capture individual cells and barcoded gel beads within droplets for reverse transcription. The libraries were prepared according to manufacturer instructions and sequenced on an Illumina NovaSeq 6000 with a paired-end 26×91 configuration targeting a depth of 50,000 reads per cell. Cell Ranger software was used to perform demultiplexing of cellular transcript data, and mapping and annotation of UMIs and transcripts for downstream data analysis.

For scRNAseq analysis, raw data was processed with Cell Ranger v6.1.0 (10x Genomics) using a standard pipeline. The downstream analysis was performed in R package Seurat v4.3.0. The following criteria was used to filter out cells: nFeatures > 200 and < 4500, % of mitochondrial genes <10, % of HBB genes < 10 and % ribosomal genes < 10. DoubletFinder v2.0.3 was used to identify and filter doublets using default parameters. The first 15 PC dimensions were used to run UMAP and the FindNeighbors functions. The clusters were identified using the FindCluster function with 0.5 resolution. Cells were annotated using SingleR v1.10.0 using the Immgen database as the reference^31^. The top 10 markers from each cluster were identified to verify SingleR cell annotation. Differential gene expression analysis was performed using the FindAllMarkers function using MAST v1.22.0 algorithm. Genes with log fold change greater than 1.3 and adjusted p-value less than 0.05 were considered up-regulated and genes with log fold less than −1.3 and adjusted p-value less than 0.05 were considered to be down-regulated genes. Gene set enrichment analysis for each annotated cell population was performed on a ranked list of all genes using fgsea v1.22.0. Mapping of gene fold changes to the KEGG oxidative phosphorylation pathway was done using Pathview v1.36.1 using default parameters. Ligand-receptor cell-cell communication predictions between cell populations were done using CellChat v1.5.0 using default parameters. Major cell populations were extracted using the subset function, and clusters within each cell population were identified by normalization and clustering as described earlier. The top 20 markers from each cluster were used to further classify and annotate the cell subsets. Differential gene expression and gene set enrichment analyses were done as described earlier. Differences in the mean between the two groups were determined using the default Wilcox test. For differential gene expression analysis and GSEA, adjusted p-values obtained from MAST and fgsea were used. Pseudotime trajectories for macrophage and monocyte subsets were constructed using R package Slingshot v2.10.089^96^. Downstream differential expression analysis was done using R package tradeSeq v1.16.090^97^. Pseudotime analysis was performed by ordering the cells and assigning resting macrophage cluster AM1 as the root node. For monocyte subsets, Mono2 subset, which was enriched in uninfected cells, was assigned as the root node. Differential gene expression analysis along the pseudotime trajectory was done using TradeSeq function fitGAM.

### Prediction of metabolic pathways using Compass

Metabolic status of cells was predicted using Compass^34^, a Python-based tool that infers flux through all reactions in a metabolic model using scRNAseq data and flux balance analysis. The Compass algorithm was run using default parameters. The output reaction penalties were transformed into reaction consistencies using a negative log where higher values correspond to higher flux through that reaction. Reactions with reaction consistencies <0.001 were removed. Using the Wilcoxon rank sum test, significant reactions were identified between the *hip1* mutant and WT Mtb. Any reaction with an FDR-adjusted p-value <0.05 was considered significant.

Differences in the effect sizes of reaction consistencies in the *hip1* mutant compared to WT Mtb were calculated using the R function Cohen’s d. The R package limma^98^ was used to calculate the fold change between *hip1* mutant and WT Mtb. Reactions were grouped by subsystem level information available from the virtual metabolic human database^99^ for downstream visualization.

### Gene Correlation Network Analysis and feature selection using LASSO

Gene correlation network analysis was performed using the R package CS-CORE^100^. Co-expression estimates and p-values were calculated using the cscore function using scaled data from Seurat as input. The top 2000 genes with the highest expression were selected for co-expression analysis. Co-expression estimates were further used as input for the R package WGCNA^101^ to calculate a dissimilarity matrix and perform hierarchical clustering using default parameters to identify modules of co-expressing genes. The moduleEigengenes function from the R package WCGNA was used to calculate eigengenes values for each module which corresponds to the module score associated with each cell, which is then segregated into treatment groups to identify modules enriched in *hip1* mutant compared to WT Mtb. The R package glmnet^102^ was used to develop a LASSO regression model. The top 2000 genes with the highest expression were selected for regression analysis. Genes with positive coefficients were selected for fitting a linear model to obtain summary statistics. Genes with p-value < 0.01 were selected as LASSO signature genes.

### Data analysis from public datasets

Mitochondrial signatures obtained from gene co-expression network analysis and LASSO were used to interrogate data from published mouse, NHP, and human transcriptomics datasets. For mouse datasets, GSE200639^50^ and GSE245950^51^ were selected. NHP datasets were obtained from two studies, GSE200151^53^ and GSE149758^52^. Human data were obtained from GSE76873^54^. Enrichment scores of MitoCore and LASSO signature genes in each dataset were calculated using gene set enrichment analysis (GSEA). Module scores were calculated using AddModuleScore function from the R package Seurat using either the MitoCore or LASSO signature genes as input. The mean module score for each sample was used for visualization.

### Metabolomics

At 2 wpi, lung tissue from individual animals was collected in a 5ml lung harvest media containing collagenase (1mg/ml) and DNase (10ng/ml). After 30 mins incubation, lungs were homogenized in a GentleMAC dissociator (Miltyeni) and centrifuged for 5 mins at 1500 rpm. The supernatant was collected as the extracellular fraction and processed for metabolomics. The cell pellet was resuspended in incomplete RPMI media containing 10% FBS and passed through a 70-μm strainer. Cells were counted, and 2-3 million cells were used for metabolomics. For processing samples for metabolomics analysis, 60 μL of the extracellular fraction (supernatant) or the cellular fraction (cell pellet) from each sample was resuspended in 205 μL acetonitrile (80%) containing internal controls (standards) in a 2 mL sterile screw cap tube. The cellular fraction from each sample was disrupted in a bullet blender machine for 1 min. The cell lysate was incubated for 30 mins at 4°C and then spun down at 14,000g for 10 mins at 4°C to precipitate any proteins, filtered through a 0.22μm filter, and stored at –80 °C. High-resolution metabolomics was performed on lung samples as described previously^65, 103^. Briefly, samples were thawed and transferred to low volume autosampler vials maintained at 4°C and analyzed in triplicate using an Orbitrap Q Exactive Mass Spectrometer (Thermo Scientific, San Jose, CA, USA) in dual positive and negative ionization mode with HILIC positive and c18 negative liquid chromatography (Higgins Analytical, Targa, Mountain View, CA, USA, 2.1 × 10 cm) using a formic acid/acetonitrile gradient. The high-resolution mass spectrometer was operated over a scan range of 85 to 1275 mass/charge (m/z) and stored as .Raw files. Data were extracted and aligned using apLCMS and xMSanalyzer with each feature defined by specific m/z value, retention time and integrated ion intensity. Three technical replicates were performed for all samples to enhance the reliability of the metabolic features measured and intensity values across replicates were median summarized. Missing data was imputed by replacing missing and zero values with ½ of the min intensity. Downstream analysis was done using the Python package Rodin^104^. Data was log-normalized, and the difference in the means between the group was assessed using one way ANOVA. Metabolite features with p-value < 0.05 were selected for PCA and heatmap visualization. Pathway analysis was conducted using mummichog, a python-based tool which predicts pathway metabolic activity based on organization of metabolic networks^105^. Pathways with p-value < 0.05 were considered significant.

### SCENITH/Puromycin quantification assay

Lung cells were isolated from WT Mtb and *hip1* mutant-infected mice at 3 wpi. 1×10^6^ cells were seeded in 96-well polypropylene plates and treated for 30 mins with each of the following metabolic inhibitors: 2-Deoxy-Glucose (2-DG, 100mM; Sigma, D8375), Oligomycin (O, 1.5 mM; Sigma, 75351), or DMSO alone (control). Following 30 min incubation with the inhibitors, puromycin (10mM; Sigma, P7255) was added to the wells and cells were incubated for another 30 mins at 37°C, 5% CO2. After treatment, cells were washed with PBS and surface-stained using fluorescently labeled antibodies. For intracellular staining, surface-stained cells were fixed and permeabilized using the Biosciences FOXP3/Transcription factor staining buffer set (Invitrogen; Cat.00-5523-00) according to manufacturer instructions. After washing, cells were incubated with PE anti-puromycin antibody (Biolegend; Cat.381584; Clone 2A4) for 30 mins at 37°C, 5% CO2. Lastly, cells were fixed in 2% paraformaldehyde (Electron Microscopy Sciences) diluted in FACS buffer and placed at 4°C until acquisition (up to 24 hr post-staining). All samples were acquired using the BD A5 Symphony using FACSDiva (BD) software and data analyzed using FlowJo software (FlowJo LLC).

### Assessment of mitochondrial mass

Isolated lung cells (1×10^6^ cells/96-well) at 3 wpi were incubated in 300nm MitoSpy Green FM probe (Biolegend) for 30 mins at 37°C, 5% CO2. Cells were washed with PBS and stained with fluorescently labeled antibodies for lung phenotyping. Stained cells were washed and fixed with 2% PFA for 2 hours. All samples were acquired by a flow cytometer BD A5 Symphony using FACSDiva (BD Biosciences) software. Mitochondrial mass was measured by Mitospy Green fluorescence (MFI) using FlowJo software (FlowJo LLC).

### ELISPOT assays

Mouse lung cells were isolated after each infection time point (week 2, 3, 8) and plated in triplicate at 1 × 10^6^ cells/well, followed by stimulation with ESAT-6_1-20_ peptide pools. Mtb-specific IL-17 and IFN-g secreting lung cells were enumerated using the ELISPOT assay. Antibodies were purchased from Mabtech (Cincinnati, OH), and the assay was performed as published previously^106^.

### Bone marrow derived macrophages (BMDM) generation

Murine bone marrow derived macrophages were generated as previously described^29^. Briefly, bone marrow cells were isolated from the femur and tibia of C57BL/6 wild type (WT) female mice under sterile condition. Isolated cells were differentiated in Dulbecco’s modified Eagle’s medium (DMEM)/F-12 medium (Lonza, Walkersville, MD) with 5% FBS, 2 mM glutamine, and 50ng/mL of recombinant mouse M-CSF-1 for 7 days on untreated tissue culture plates at 37°C with 5% CO2. Cells were replenished with freshly prepared media on days 3 and 5. On day 7 cells were counted and seeded onto treated tissue culture plates for experiments.

### Transmission electron microscopy and mitochondrial morphology measurements

BMDMs were infected with *hip1* mutant or WT Mtb at MOI=3 for 24 hours. Macrophage monolayers were fixed while attached to culture plates *in situ* using an aqueous 2% paraformaldehyde solution, removed from the BSL3 lab, and subsequently washed in dH_2_0. The monolayers were then further fixed using a mixture of 2.5% glutaraldehyde, 1.0% paraformaldehyde, 2.6mM MgCl_2_, 2.6mM CaCl_2_, 50mM KCl, 0.01% picric acid, and 2% sucrose in 0.1M cacodylate buffer. Cells were then post-fixed in 1.0% osmium tetroxide for 1 hour, washed twice with deionized water (10 minutes each), and then stained *en bloc* with 2% aqueous uranyl acetate in deionized water for 20 minutes at 60 ^°^C. Monolayers were next dehydrated in a stepwise manner in 25% ethanol containing 1.0% p-phenylenediamine, 50% ethanol, 70% ethanol containing 0.5% tannic acid, and 95% ethanol, followed by three changes in 100% ethanol. Each dehydration step was carried out for 10 minutes. Cells still attached to culture plates were infiltrated with Eponate 12 resin according to the manufacturer’s instructions and polymerized for 48 hours at 60 ^°^C. Polymerized blocks containing monolayers flat-embedded in the bottom of the blocks were then separated from the culture plate. Ultrathin sections were cut from the bottom of the blocks with a Leica EM UC6 ultramicrotome, stained with uranyl acetate and lead citrate, and imaged in a JEOL JEM 1400 TEM operated at 80KV. Electron micrographs were acquired on a 2048×2048 charge-coupled device (CCD) camera (UltraScan 1000, Gatan Inc, Pleasanton, CA, USA). The Blend Montages option in Etomo (a tomographic reconstruction program in the IMOD suite) was used to create a blended stack to be used for data analysis ^107^. The images were processed using Fiji software. Each mitochondrion in the image was manually marked using the polygon tool to generate a region of interest (ROI). For each ROI, parameters such as area, circularity, roundness, and aspect ratio were measured using the measure tool. The data for each ROI was saved as a csv file and plotted in R Studio software.

### Quantification of ATP concentrations

BMDM were infected with either the WT Mtb or *hip1* mutant, followed by treatment with the metabolic inhibitor, Oligomycin (6mM; Sigma, 75351) for 24 hours or with no inhibitor as controls. After 24 hours, the cell lysate was collected and filtered (0.2 microns). The levels of ATP from the cell lysate were measured using the ELISA Kit (Abcam, ab83355) according to the manufacturer’s instructions.

### Statistical analysis

Statistical analyses of data were conducted using R-studio (v4.3.2) and GraphPad Prism (v10.6.1). Two-sided Wilcoxon rank sum test, two-sided unpaired student t-test and one-way ANOVA test with false discovery rate (FDR) for multiple comparison corrections was used for statistical analysis. Statistical tests performed for each figure are noted in each figure’s legend. The model figures were generated using BioRender.com.

## Data Availability

All raw and processed scRNAseq data will be deposited in GEO and will be available upon acceptance of the manuscript. All the data used to generate figures are provided via the GitHub repository described in “Code availability.” All data are included in the Supplementary Information or available from the authors, as are unique reagents used in this article.

## Code Availability

All R and python scripts used for the analysis and to figure generation are available in the GitHub repository at https://github.com/prashantbajpai/Mouse_2week_transcriptomics.

---

**Extended Data Fig. 1.**
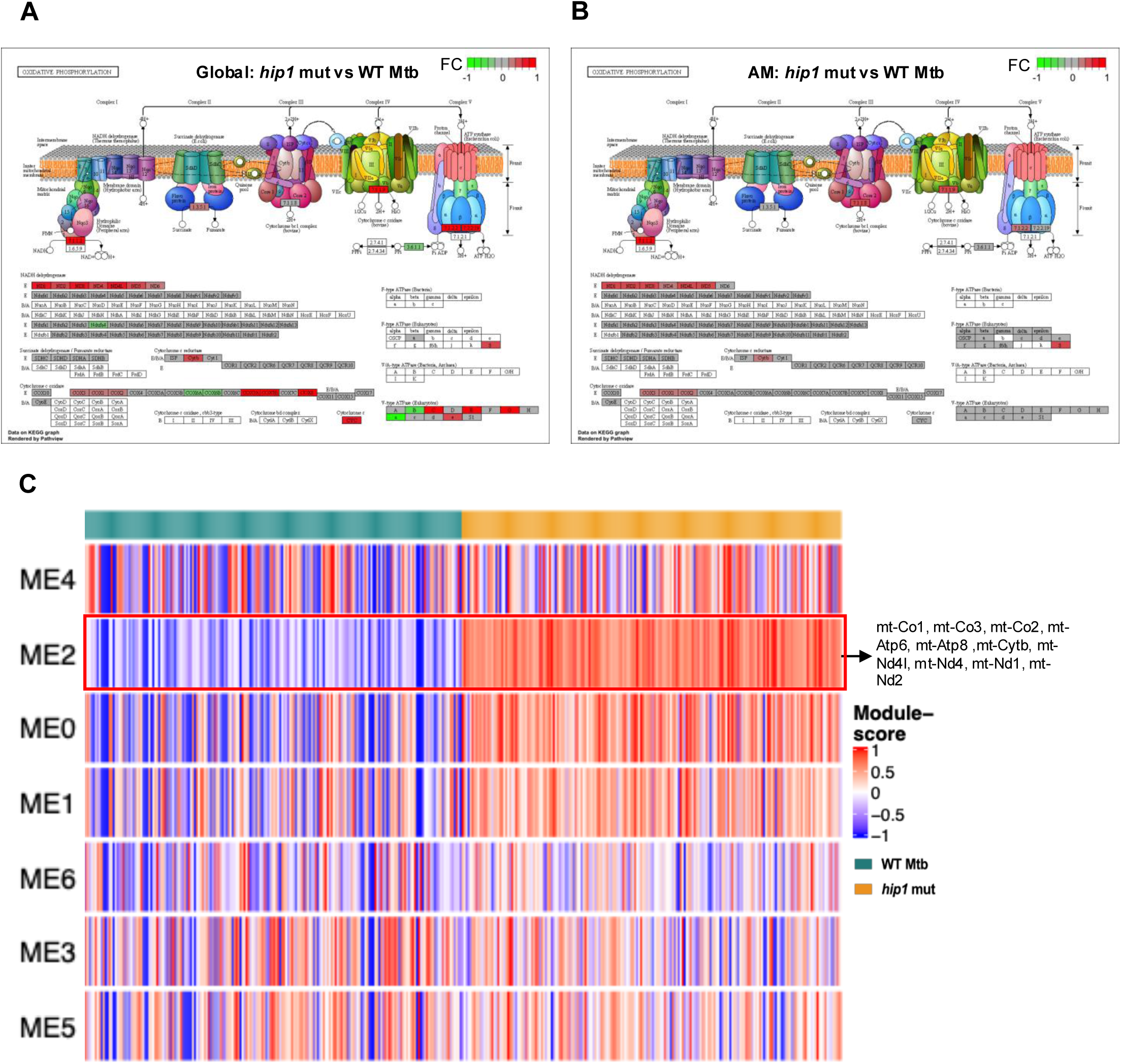
KEGG Pathview of total and AM mitochondrial ETC. **a,b**, Representation of DEGs superimposed on mitochondrial OXPHOS, ETC pathways (KEGG) using Pathview. Genes are colored by log_2_ fold change between the *hip1* mutant vs WT Mtb. Genes with greater fold change in *hip1* mutant compared to WT Mtb are shown in red, while genes with greater fold change in WT Mtb compared to *hip1* mutant are shown in green. **(a)** Global KEGG Pathview with all cells combined, **(b)** KEGG Pathview of AMs. **c**, Heatmap shows the correlation between genes from clusters obtained by gene correlation network analysis and outcome group (WT Mtb or *hip1* mutant). Each row shows the different clusters, whereas the column represents genes from each cluster. The heatmap is colored by correlation value from low (blue) to high (red) correlation.

**Extended Data Fig. 2.**
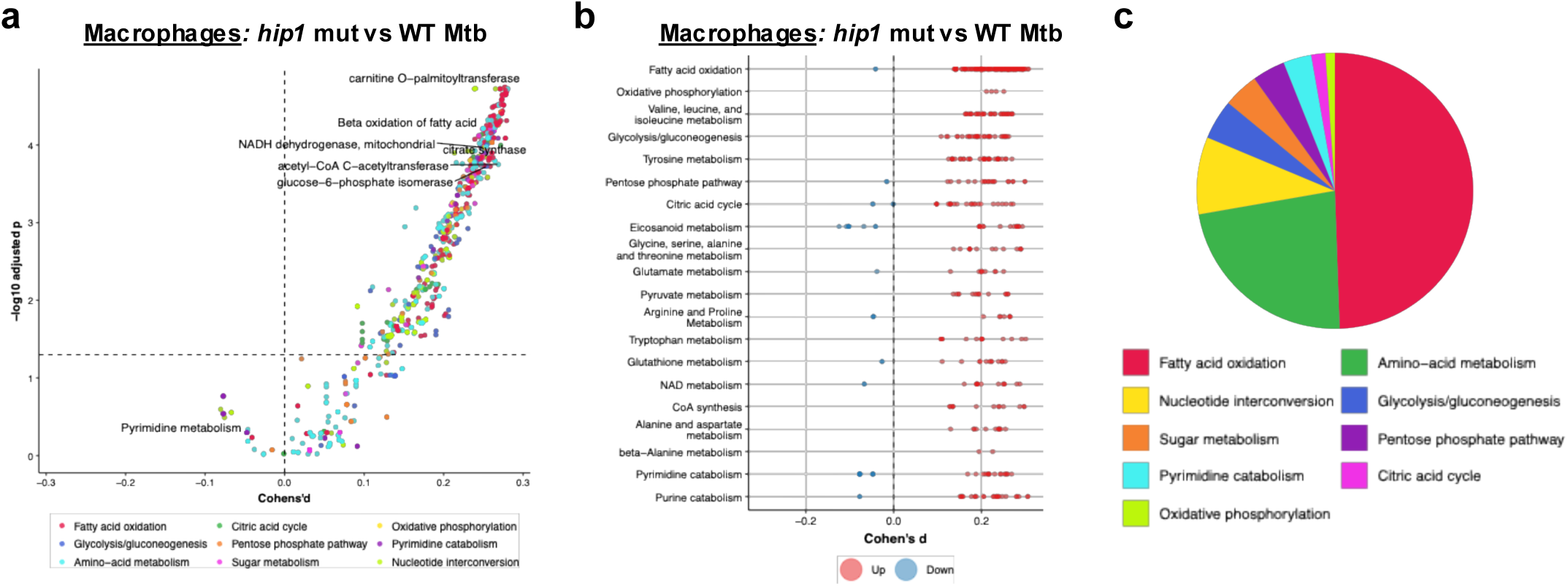
Prediction of metabolic signatures in lung macrophages using Compass. **a**, Differential metabolic activity assessments between the *hip1* mutant *versus* WT Mtb based on the Compass^34^ score for each metabolic reaction, predicting multiple metabolic processes enriched in *hip1* mutant-infected compared to WT Mtb. The effect size was assessed using Cohen’s d statistic, shown on the x-axis, calculated as the difference between sample means over the pooled sample standard deviation. Probability estimates are derived from Wilcoxson’s rank sum test shown in y-axis. Each dot represents one metabolic reaction colored by reaction subsystem (shown in c). **b**, Representation of macrophage metabolic reactions (listed on y axis) comparing *hip1* mutant vs WT Mtb using Compass, where each reaction (dot) is partitioned by Recon2 pathway^108^ and colored by the sign of their Cohen’s d statistic value. Reactions with positive Cohen’s d (significantly enriched in *hip1* mutant) are shown in red, while negative Cohen’s d (enriched in WT Mtb) are shown in blue. **c**, Pie chart depicting the distribution of macrophages metabolic reactions grouped by Recon2 pathways with Cohen’s d > 0 when comparing *hip1* mutant *versus* WT Mtb. Individual reaction categories are labeled by the different colors shown.

**Extended Data Fig. 3.**
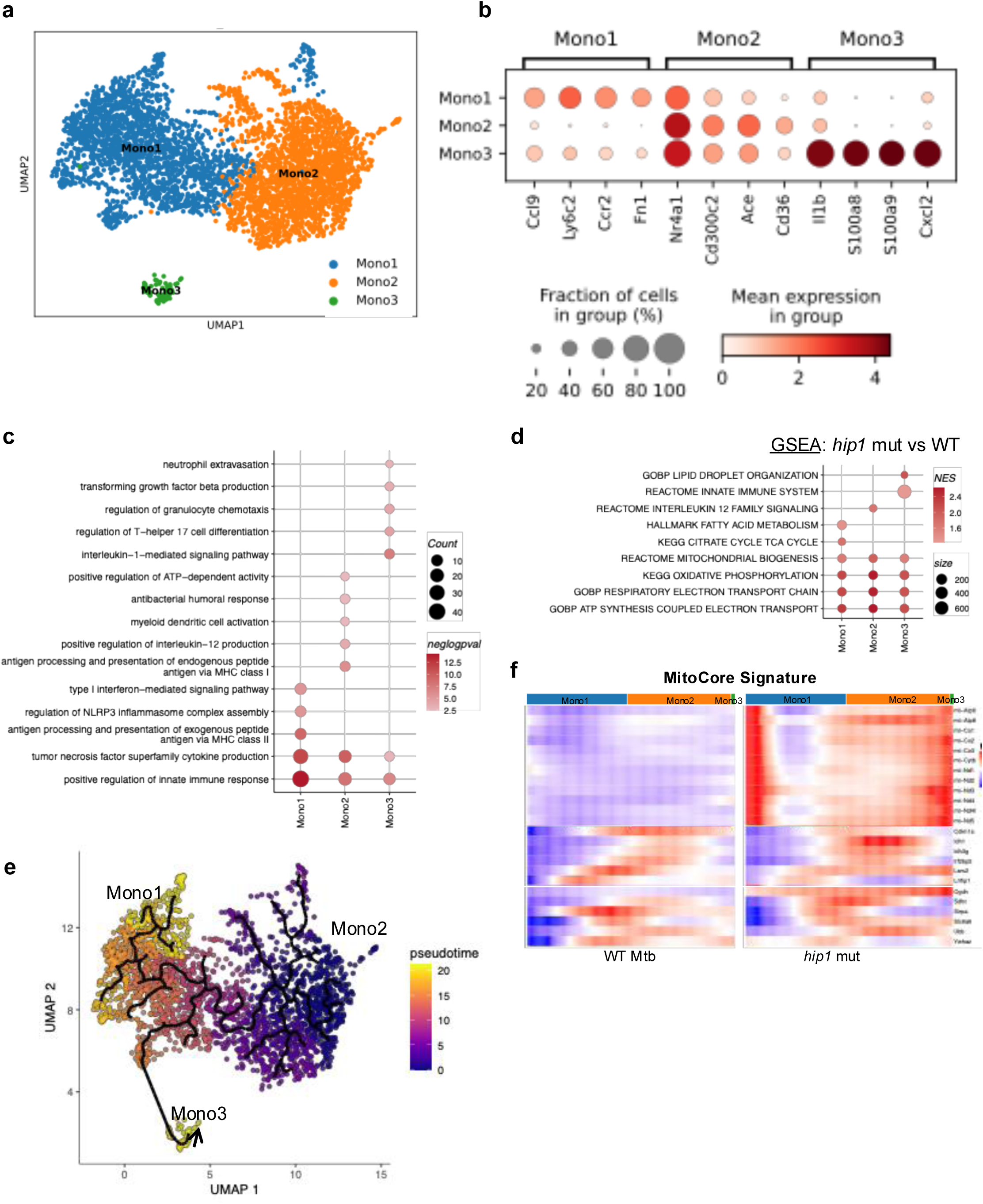
Phenotypic and functional signatures associated with lung monocyte subsets. **a,** UMAP projection of concatenated monocytes from all three groups (n=3 per group) segmented into Mono1, Mono2, and Mono3 subsets based on Seurat clustering using the nearest neighbor algorithm. **b**, Bubble dot plot shows expression of the key genes that classify each monocyte subset into Mono1, Mono2, and Mono3. The size of the dot indicates the fraction of cells expressing the gene, and the intensity of the color reflects the extent of expression of the gene. **c**, Bubble plot shows pathways differentially enriched in each monocyte subset compared to others. For each subset, genes with a fold change > 1.3 and FDR-adjusted p-value < 0.05 were selected for pathway analysis. Bubbles are colored based on p-value, where a darker color represents a lower p-value. The size of each bubble represents the number of genes in the reference pathway. **d**, Gene set enrichment analysis (GSEA) showing the enriched pathways comparing *hip1* mutant versus WT Mtb monocyte subsets. Pathways with p-value < 0.05 obtained from GSEA analysis were selected for visualization. The bubble plot shows the upregulated (red) or downregulated (blue) pathways for each subset, whereas the size of the bubble represents the number of genes in the reference pathways. **e**, Combined UMAP projection of *hip1* mutant and WT Mtb showing monocyte subsets in a pseudotime scale. The color represents the direction from early (violet) to later (yellow) stages in pseudotime, and arrows show different pseudotime trajectories. **f**, Heatmaps depicting the mitochondrial core gene signature in monocyte subsets on a pseudotime scale from Mono1 to Mono2 to Mono3. Each row represents a gene with the color gradient indicating the expression level (red for high, blue for low).

