## Supplementary figures and tables for "Multimodal profiling reveals *Mycobacterium tuberculosis* restricts lung mitochondrial immunometabolism to promote pathogenesis"

#### Supplementary Fig.1|

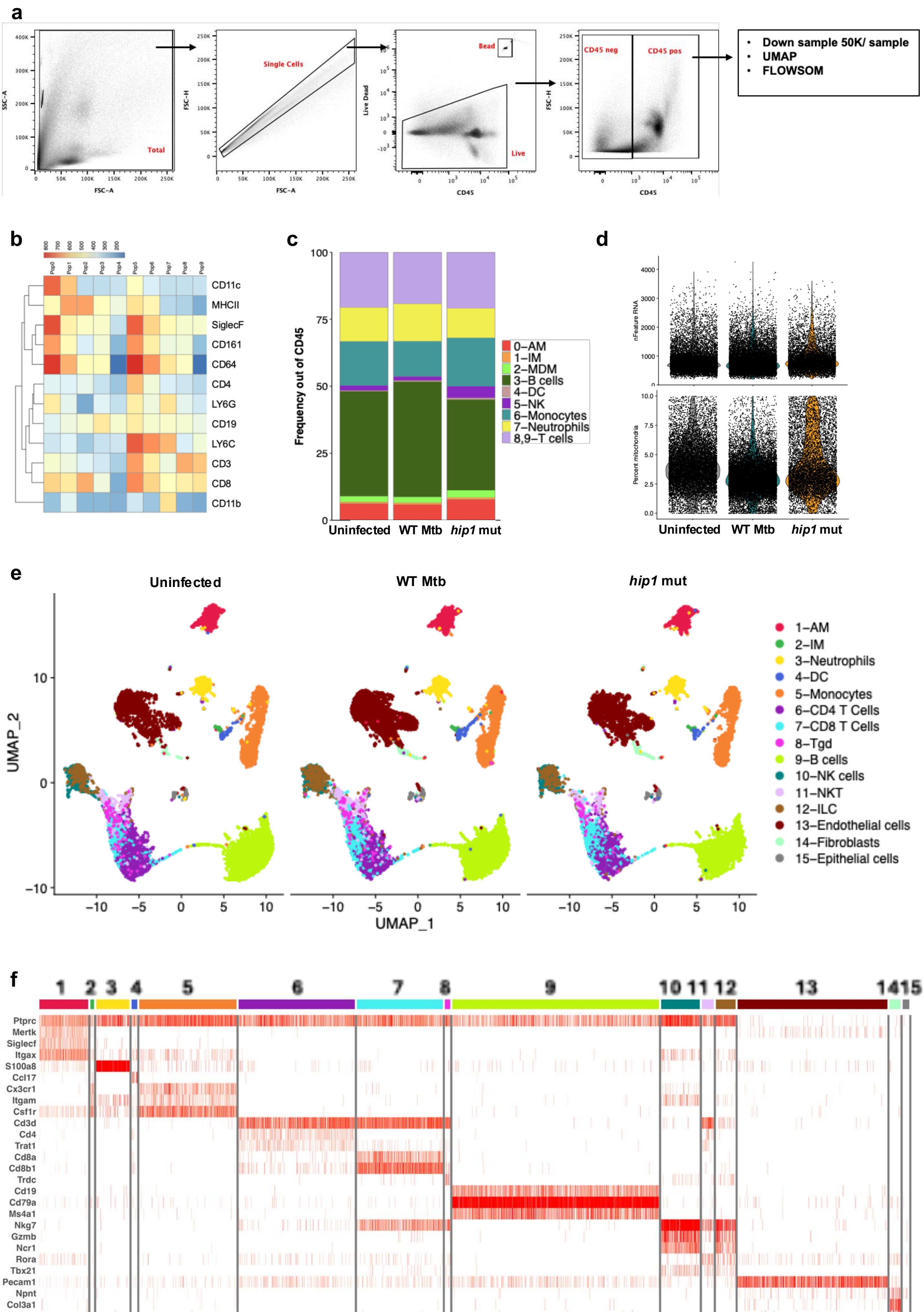

Supplementary Fig.2|

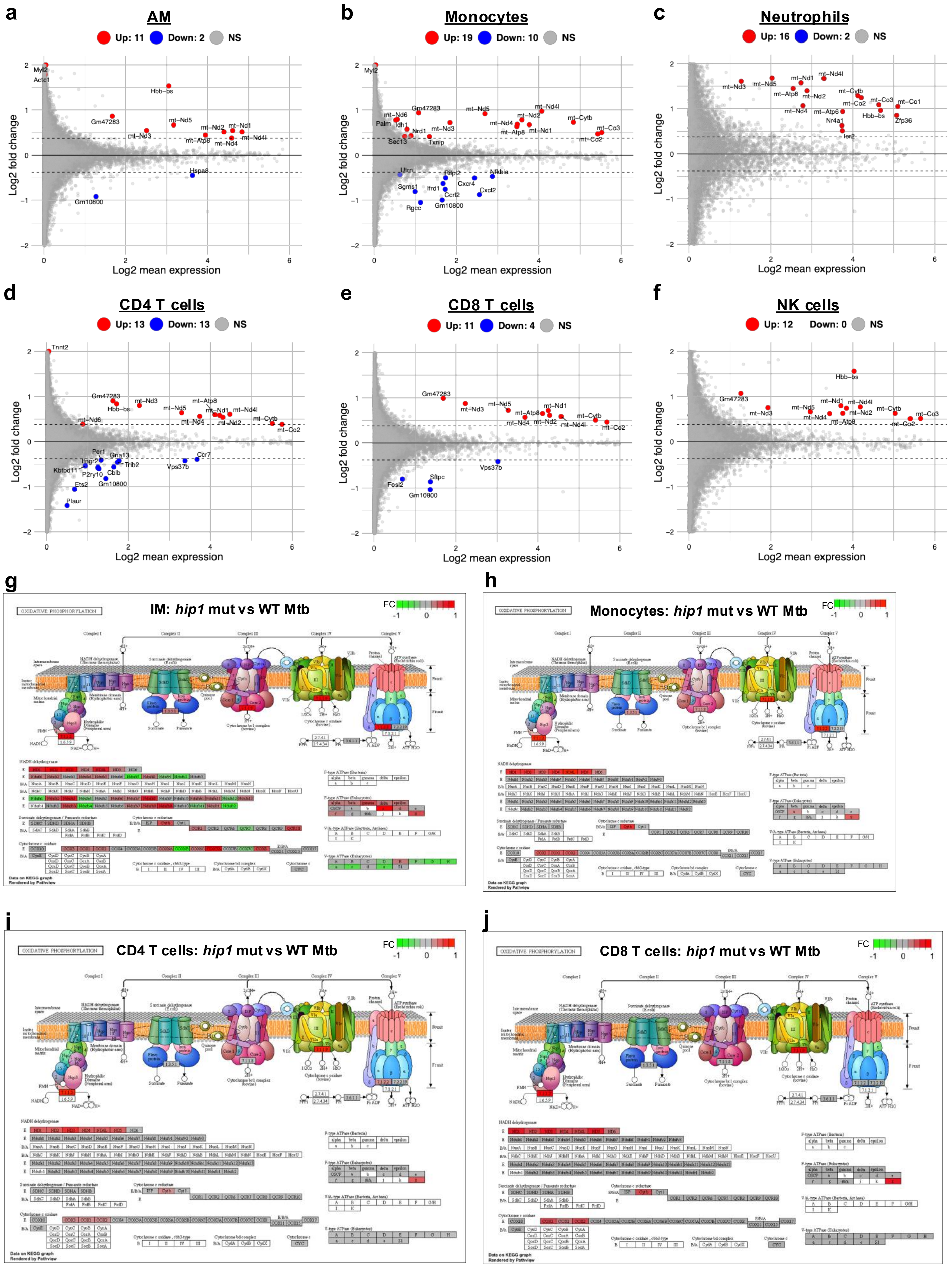

**Supplementary Fig.3|**

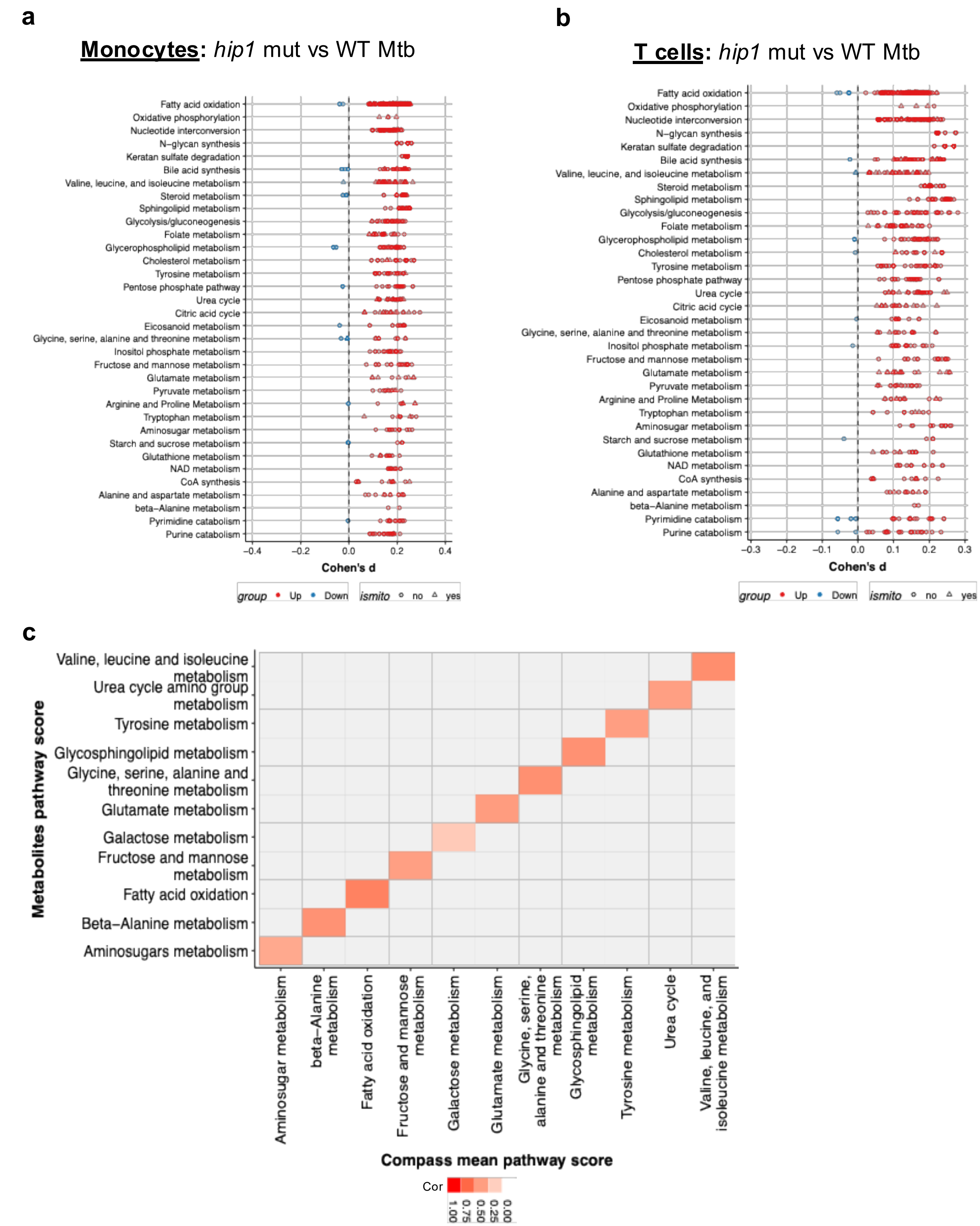

**Supplementary Fig.4|**

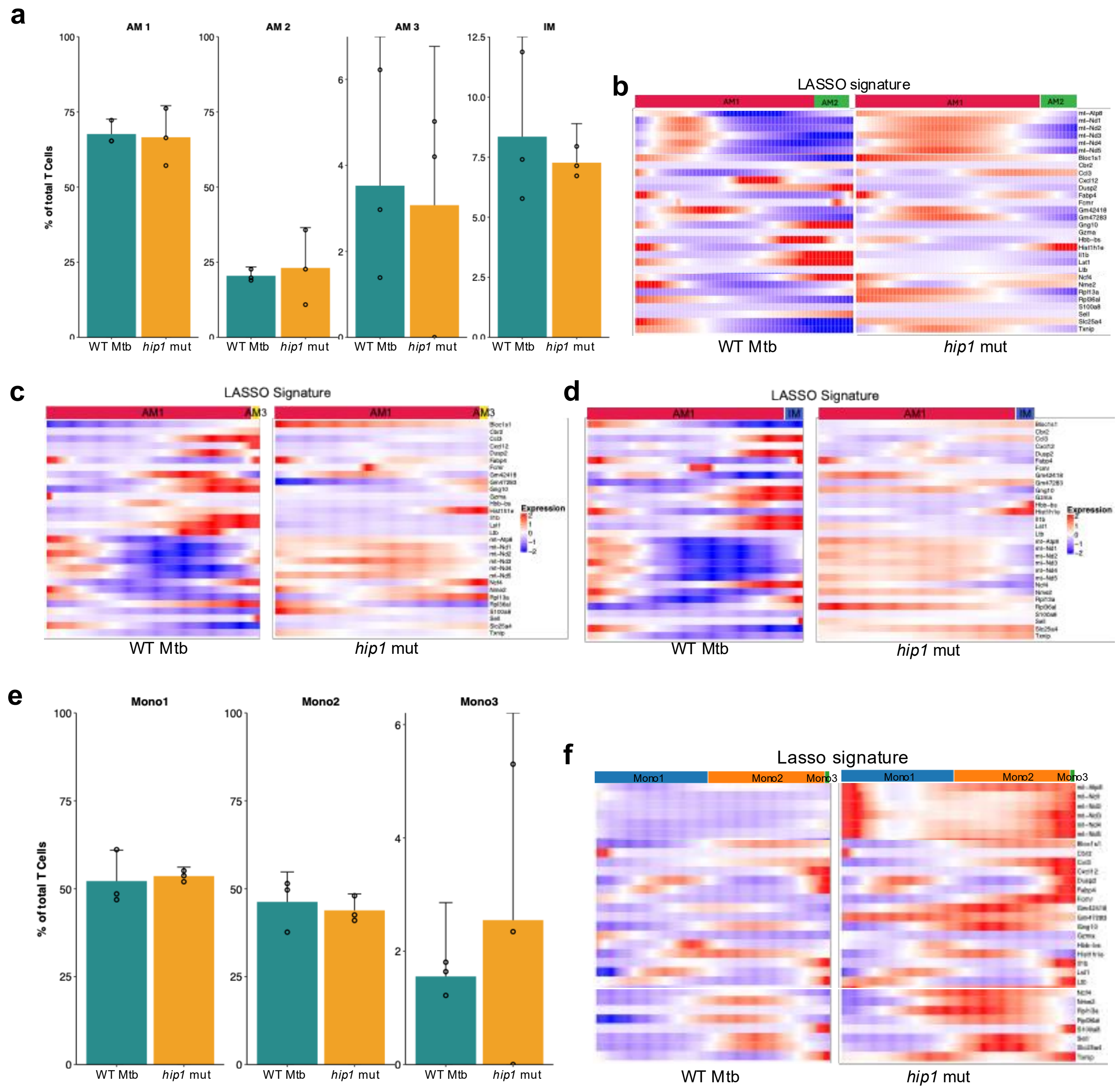

Supplementary Fig.5|

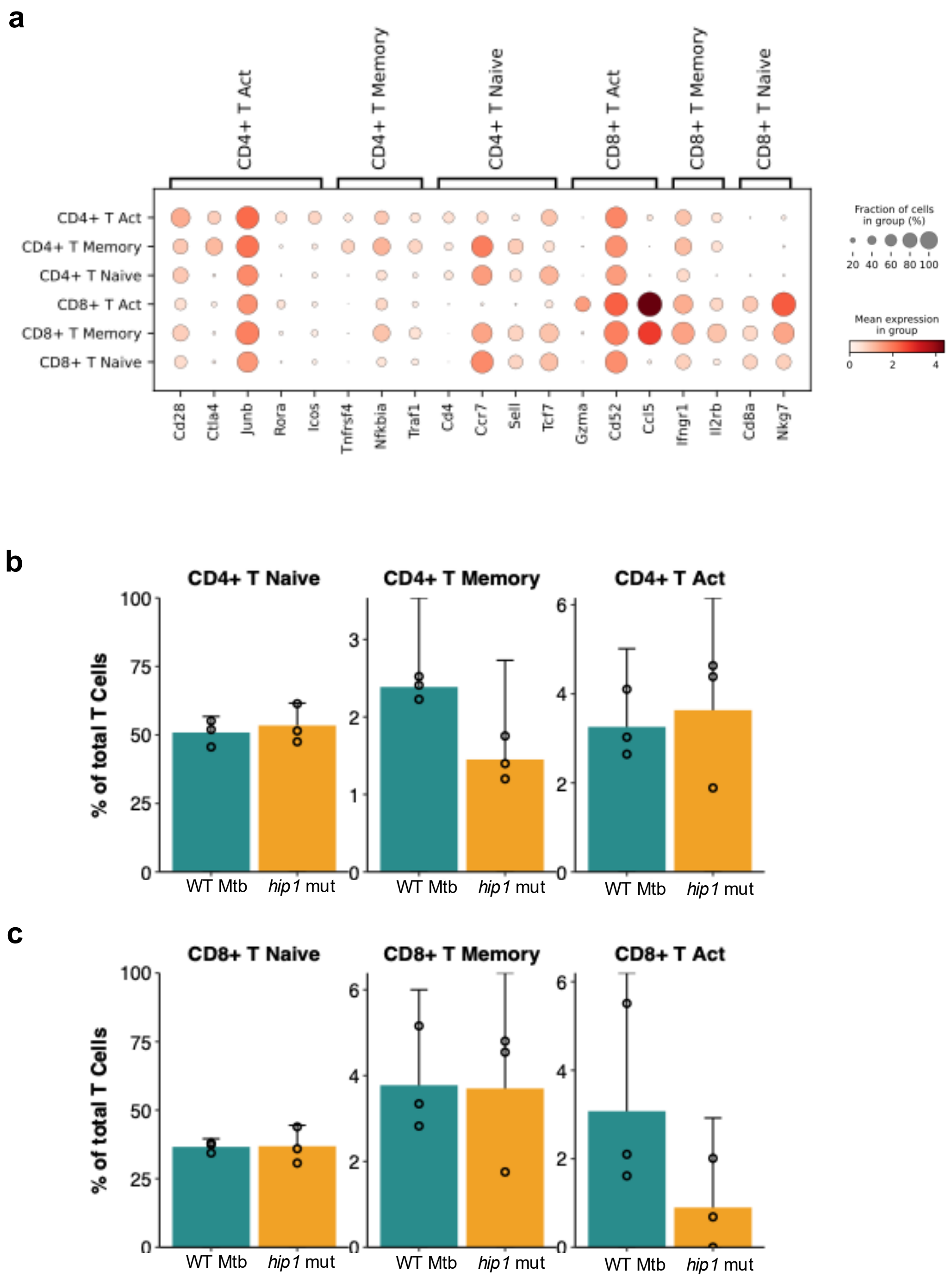

**Supplementary Fig.6|**

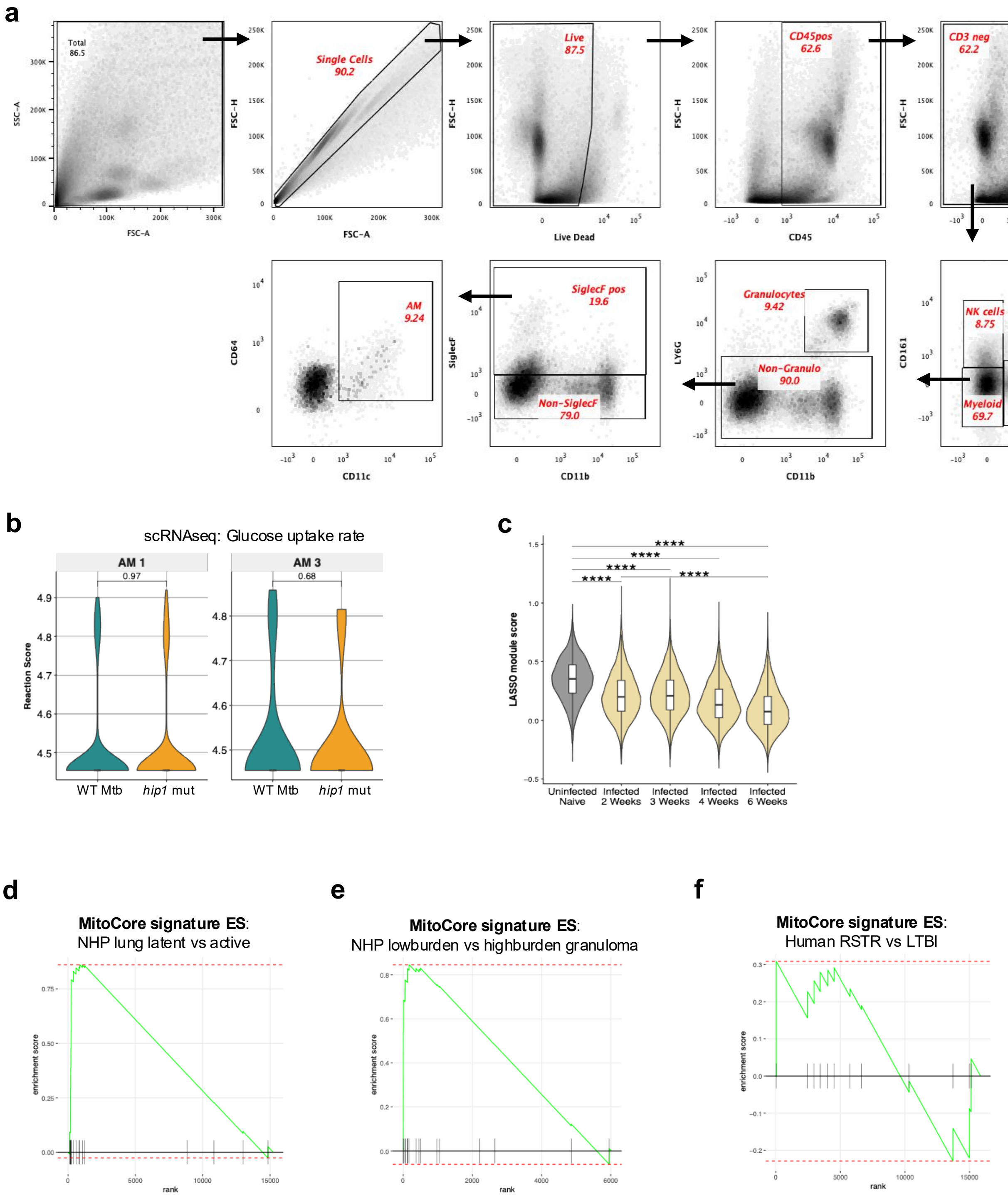

### Supplementary Table 1|

| SampleID | Group | AM | IM | Neutrophils | DC | Monocytes | CD4 T Cells | CD8 T Cells | Tgd | B cells | NK cells | NKT | ILC | Endothelial cells | Fibroblasts | Epithelial cells |
|---|---|---|---|---|---|---|---|---|---|---|---|---|---|---|---|---|
| 1 | Uninfected | 321 | 55 | 370 | 84 | 1045 | 735 | 668 | 34 | 2068 | 364 | 86 | 194 | 1061 | 56 | 51 |
| 2 | Uninfected | 324 | 3 | 23 | 2 | 163 | 555 | 315 | 12 | 550 | 65 | 16 | 13 | 66 | 4 | 1 |
| 3 | Uninfected | 313 | 1 | 59 | 3 | 314 | 885 | 519 | 46 | 990 | 102 | 41 | 26 | 5 | 1 | 2 |
| 4 | WT Mtb | 212 | 13 | 153 | 20 | 443 | 567 | 424 | 28 | 816 | 244 | 68 | 127 | 1529 | 104 | 52 |
| 5 | WT Mtb | 267 | 36 | 241 | 27 | 610 | 768 | 518 | 28 | 1188 | 252 | 105 | 128 | 1485 | 117 | 61 |
| 6 | WT Mtb | 200 | 16 | 187 | 52 | 656 | 439 | 414 | 25 | 1351 | 181 | 72 | 162 | 843 | 60 | 40 |
| 7 | *hip1* mut | 13 | 1 | 35 | 8 | 50 | 75 | 39 | 3 | 131 | 28 | 4 | 8 | 95 | 18 | 7 |
| 8 | *hip1* mut | 220 | 19 | 74 | 16 | 283 | 295 | 288 | 14 | 406 | 124 | 28 | 43 | 301 | 28 | 18 |
| 9 | *hip1* mut | 222 | 16 | 281 | 31 | 597 | 656 | 488 | 40 | 1347 | 296 | 90 | 139 | 1026 | 87 | 48 |

**Supplementary Table 2|**

| **Name** | **Dataset type** | **Source** | **Reference** |
|---|---|---|---|
| Mouse Mtb infection 2, 3,4, 6 weeks | ScRNAseq | Mouse lung | Pisu, D., *et al. Nat Commun* 15, 8522 (2024). |
| Mouse Mtb infection day 50, day 100. | ScRNAseq | Mouse lung | Akter S et.al. Cell Rep. 2022 Jun 21; 39(12):110983. |
| Human CD14 monocytes from RSTR and LTBI | Microarray | Human PBMC | Seshadri C et.al. PLoS One. 2017 Apr 17;12(4):e0175844. |
| High and low burden granuloma 10 weeks post Mtb infection | ScRNAseq | NHP lung | Gideon HP et.al. Immunity. 2022 May 10;55(5):827-846.e10. |
| Active TB and LTBI | ScRNAseq | NHP lung | Esaulova E et.al. Cell Host Microbe. 2021 Feb 10;29(2):165-178.e8. |
